# The interplay of sexual selection and hybridization can drive sexual radiation

**DOI:** 10.1101/2025.01.28.635380

**Authors:** Kotaro Kagawa

**Affiliations:** Ecological Genetics Laboratory, National Institute of Genetics, Mishima, Shizuoka, Japan; Graduate School of Life Sciences, Tohoku University, Sendai, Japan

## Abstract

Sexual selection is considered a major driver of species diversification, but little is known about whether and how sexual selection may contribute to the incipient stages of speciation and sexual radiation. Based on evolutionary simulations, here I propose that an interplay of sexual selection and hybridization can drive the rapid formation of multiple species with diverse exaggerated sexual displays within a lineage. Existing theories suggest that sexual selection can cause evolutionary dynamics with multiple stable equilibria, with which multiple species with distinct sexual displays and mate preferences can persist stably. However, the stability of each equilibrium hinders the formation of new species from an existing species that is already occupying one of the stable equilibria. In such situations, hybridization between phenotypically similar but genetically distinct lineages can catalyze speciation. Hybridization can generate genetic variation by recombining alleles from different lineages. I provide evolutionary simulations demonstrating that hybridization that gives rise to genetic variation in mate preference can modify the sexual selection regime to open up opportunities for the evolution of sexually selected displays toward previously unoccupied stable equilibria. Hybridization thus can trigger evolutionary shifts between alternative evolutionary equilibria that sexual selection generates, driving the incipient formation of sexual radiation.

## Introduction

Male sexual displays for attracting mates such as nuptial coloration, mating song, and courtship dance often exhibit spectacular elaboration and interspecific diversity. Diversification of sexual displays and mate preference within a lineage can promote incipient stages of evolutionary radiation because interspecific differentiation in these traits can cause behavioral reproductive isolation [1–4]. In line with this idea, young evolutionary radiations without complete postzygotic and geographic isolation between species often involve tremendous interspecific variation in male secondary sexual displays [5–10]. Interestingly, some of such evolutionary radiations consist of ecologically similar species (i.e. sexual radiation) [6,7,11–13]. In sexual radiations, non-ecological speciation via evolutionary diversification of mate preference and sexual displays may serve as the primary driver of speciation.

Inter-sexual selection, which typically occurs through female mate choice, can drive the evolution of costly exaggerated sexual displays and is also considered a major driver of diversification of sexual displays [1–3]. In support of this view, several theoretical models of inter-sexual selection revealed that evolutionary dynamics of mate preference and sexual displays can have multiple alternative stable equilibria in which different exaggerated phenotypes of sexual displays are stably maintained [14–18]. In each stable equilibrium, female mate preference causes sexual selection for an exaggerated sexual display thereby offsetting the natural selection against costly sexual displays. Multiple alternative stable equilibria have been identified in several previous evolutionary models considering various different mechanisms for the evolution and maintenance of female mate preference including “direct benefits” of mate choice [15] and indirect genetic benefits via “good gene” effect [14,15,17] as well as “sexy son” effect (or Fisherian selection) [16]. Therefore, sexual selection generally can set up conditions where there are multiple alternative stable equilibria of evolution, which can explain the evolutionary persistence of interspecific diversity of mate preference and sexual displays.

However, the presence of alternative stable equilibria is not sufficient for explaining the incipient process of speciation. As shown by a previous theoretical study [17], it is true that multiple species occupying different evolutionary equilibria can be formed if evolution is started from populations of an ancestral species with an evolutionarily unstable combination of mate preference and sexual displays. However, such an initial state is unlikely to be common in nature since evolutionarily unstable phenotypes are ephemeral. Importantly, once a species reaches one of evolutionary stable equilibria, evolution to deviate from the equilibrium will be suppressed. Therefore, the evolution of novel mating trait phenotypes, which is necessary for the inter-population diversification of mate preference and sexual displays, is unlikely to occur in a species that has already reached one of stable equilibria. Thus, it remains unclear whether and how sexual selection can contribute to the incipient stages of speciation.

There is a growing recognition that genetic variation arising from hybridization can promote rapid evolution and adaptive radiation [6,19–21]. In hybrid populations, segregation and recombination of genetic materials from different parental lineages can create diverse novel genotypes. Hybrid genotypes can generate various phenotypes including not only intermediates between parental forms but also transgressive segregants exceeding phenotypic ranges of parental lineages combined, which can facilitate adaptation to previously unoccupied ecological niches [6,19–21]. In addition to promoting ecological speciation, hybridization may also promote non-ecological speciation by generating novel phenotypes of mating traits, including sexual displays [22]. In line with this idea, there are some known cases of hybrid speciation with diversification of sexual display [23] as well as hybridization causing transgressive segregation of male sexual displays [24–28]. Additionally, genetic evidence of past hybridization events has been detected in many sexual radiations and adaptive radiations with diversification of sexual displays [12,13,29–36]. However, it remains unclear whether and how new species with novel exaggerated sexual displays may be formed from a hybrid population.

Here, I propose that an interplay of hybridization and sexual selection on multiple sexual displays can promote incipient formation of sexual radiation through evolution of diverse exaggerated sexual displays. This hypothesis considers sexual selection causing multiple alternative stable equilibria of evolution and hybridization between two genetically diverged populations within the same species. The two populations are descendants of the same common ancestral population that has already reached one of alternative stable equilibria. The ancestral phenotypes will be maintained in both populations since they constitute a stable equilibrium of evolution. However, the genetic basis of the same phenotypes may diverge between two populations through genetic drift if there are many alternative genotypes generating the same phenotype; such a many-to-one relationship between genotype and phenotype is common for polygenic traits [37]. Hybridization thus can generate transgressive phenotypic variation in both female mate preference and male sexual displays through reshuffling of alleles inherited from different parental lineages with similar phenotypes. In the parental populations, a strong directional sexual selection for an exaggerated male display is generated because of the concordance of mate preference among females, and the sexual selection offsets the natural selection against the costly display to maintain the exaggerated male display. In the hybrid population, on the other hand, the temporal increase of genetic variation in female mate preference will weaken the directional sexual selection. Hence, hybridization can lead to the evolutionary loss of the costly male display. Subsequently, if genetic variation in female mate preference drops in the hybrid population, sexual selection can drive the evolution toward one of alternative stable equilibria again, most of which have not been occupied by the ancestral and parental lineages. This process can form a new species with a novel exaggerated sexual display that is reproductively isolated from the parental lineages. Furthermore, if there are many alternative stable equilibria of evolutionary dynamics in the phenotype space of multiple male sexual display traits, recurrent episodes of hybridization between the same pair of parental lineages may yield many new species that occupy different evolutionary equilibria, forming a sexual radiation with diverse exaggerated sexual displays.

To examine the theoretical validity of this hypothesis, I conduct individual-based simulations considering hybridization and sexual selection on multiple male sexual display traits. Sexual selection driving the evolution of exaggerated male sexual displays requires mechanisms for the evolution and maintenance of biased female mate preference. As such, the simulation model considers the “direct benefit” of mate choice [38,39], which is one of the most prevalent mechanisms for the maintenance of female mate preference for costly male displays. The direct benefit model posits that biased female mate preference can evolve and persist when female choosing mates with specific phenotypes can gain direct fitness benefits such as greater egg fertilization rates, access to better breeding sites, avoidance of parasites transmission from mates, and better paternal care to her offspring [40–42]. Especially, the evolution and maintenance of female mate preference for exaggerated male displays can be driven if females gain direct benefits by choosing males in good conditions and if the exaggerated male displays honestly reflect the condition of males [38,39]. Evidence from many animal species supports that males in bad conditions (e.g. infected with parasites) often cannot express or maintain exaggerated sexual displays, supporting that exaggerated sexual displays can honestly signal male conditions [41,43–45]. In a previous model considering such a direct benefit of mate choice [15], there are multiple alternative stable equilibria of evolution; in different equilibria, different exaggerated male display phenotypes signal the male conditions. In the present study, I first explore conditions for the existence of alternative stable equilibria in a model considering a direct benefit of mate choice. Then, I simulate evolutionary dynamics with hybridization under a condition where there are multiple stable evolutionary equilibria. The simulations consider two scenarios for hybridization: (1) hybridization in a secondary contact zone between two populations and (2) hybridization caused by a cycle of temporal isolation and reconnection of four populations.

## The model

### Male traits and the benefit of mate choice

The individual-based model (implemented in Java) simulates evolution of male secondary sexual display and female mate preference in sexually reproducing diploid organisms. Male sexual display consists of two quantitative traits. Phenotype of each sexual display depends on both genotype for *L* genetic loci and another non-genetic male trait, the resource budget. Resource budget represents the amount of resources (e.g., energy and time) available for reproductive efforts including the expression of sexual display, courtship, sperm production, territory guarding, and parental care. For simplicity of discussion, I hereafter refer to these male investments for increasing the mate’s reproductive success, egg fertilization rate, and offspring fitness collectively as paternal investment. The model assumes that the resource budget of each male individual *i*, *E_i_*, is subject to environmental variance; *E_i_* is assigned as a random value drawn from the normal distribution *N*(*E_max_*/2, *_e_*), while if the random value is smaller than 0 or larger than *E_max_*, *E_i_* is set to 0 or *E_max_*, respectively. Males express two sexual displays by allocating available resources to them, which tradeoffs with their paternal investment. Phenotypes of two display traits of a male individual *i*, *D_si_* (*s* = 1, 2), are given as: *D_si_ = d_si_ E_i_* /(1+Σ*_s_*|*d_si_*|), where *d_si_* is a genetic trait that determines the relative resource allocation to the *s*-th display trait. Negative values of *d_si_* give negative values for the display phenotype *D_si_*, and *D_si_* = 0 represents the optimal phenotype with the minimal resource demand, which in turn maximizes the paternal investment *f_i_*. The resource allocation to paternal investment is fixed to 1; thus, *f_i_* = *E_i_* /(1+Σ*_s_*|*d_si_*|).

A possible interpretation of this model is that *D_si_* represents the expression level of a pigment involved in nuptial coloration, where *D_si_* = 0 corresponds to the most cryptic coloration. Positive and negative values of *D_si_* thus correspond to overexpression and underexpression of the pigment compared to the most cryptic coloration, respectively. In this case, the cryptic coloration would help to minimize the energy for hunting and/ or avoiding attacks by predators as well as same-sex rivals in intrasexual competition; conversely, both overexpression and underexpression of the pigment demand extra energy.

The model assumes that the reproductive success of a mating pair increases with the paternal investment by the male partner. Therefore, females can gain direct fitness benefits by choosing males with high *f_i_*. At the same time, both paternal investment *f_i_* and the degree of display exaggeration (Σ*_s_*|*D_si_*|) increases with the resource budget, *E_i_*, unless *d*_1*i*_ and *d*_2*i*_ are zero. This means that females can indirectly choose males with higher *f_i_* by choosing males with more exaggerated sexual displays. However, the positive correlation between *f_i_* and (Σ*_s_*| *D_si_*|) is weakened when genetic variation in *d*_1*i*_ and *d*_2*i*_ is high among males.

### Female mate preference

Female mate preference can be described as a “preference function” that maps female response (e.g. the probability of acceptance) to every possible male phenotype [46]. The model assumes that each female individual *j* has two preference functions, each of which assigns preference scores, *P_sij_*, for the *s*-th sexual display (*s* = 1, 2) of every male individual *i*. The shape of the preference function changes depending on female genotype. To evaluate the robustness of simulation results to changes in the way to model the preference function, I conduct simulations with three alternative models of preference function: the exponential-, Gaussian-, and beta preference function models (Fig. S1). As simulation results have suggested that the effect of hybridization on evolutionary dynamics are qualitatively similar under all three models, the main text focuses on results with the beta preference function model, and results with other two models are shown in supplementary files.

The beta preference function model uses the beta distribution function to model the mate preference function. The beta distribution function is useful for modeling mate preference [47] because it can represent diverse shapes with only two positive parameters *α* and *β* (Fig. S1c). Since females have separate preference functions for each male display, there are four female-specific quantitative traits, *a_s_* and *b_s_* (*s* = 1, 2). Four positive parameters of two beta distribution functions are given as: *α_s_* = *exp*(*a_s_*) and *β_s_* = *exp*(*b_s_*). Supplementary text S1 provides details of the preference function models.

### Genetic basis of traits

The model assumes that all genetic traits are quantitative and controlled by independent sets of *L* genetic loci located at random positions of the genome with 15 autosomal chromosomes of 200cM long (there is no sex chromosome). All loci additively affect the traits, and there is no epistasis and pleiotropy. Point mutation within an allele of a locus changes the phenotypic effect of the allele by adding a value drawn from a normal distribution, *N*(0, *_m_*). Each locus is 0.005cM long, and the position of a mutation within the locus is randomly assigned when it occurs. Effect of the allele carrying no mutation is set to 0. Point mutation and crossover between chromosomes occur at rates *μ*/locus and 0.01/cM, respectively, when the genome of a new individual is generated from parental genomes through meiosis.

### Mating and reproduction

The model assumes discrete generations without overlapping. Each generation consists of three phases: mate choice, reproduction, and density regulation in the juvenile stage. In the mate choice phase, each female individual has up to *E* times of opportunities to encounter a male individual randomly drawn from the population. *E* equals 1/*θ*, where *θ* is the cost of searching for the next male. The probability that the female individual *j* accepts the male individual *i* upon an encounter is given by the product of preference scores for two display traits, Π*_s_P_sij_*. This assumption best fits the case where females evaluate two sexual displays of a male independently at different stages of courtship to decide whether or not to reject him. Females that have rejected all males they encountered fail to reproduce. Females can mate only once, whereas males can mate with up to *M* females. Then, the reproduction phase takes place, during which each mating pair produces offspring. The number of offspring depends on both male’s paternal investment (*fi*) and mate search cost for the female. The expected number of offspring, λ, is given as: λ = 100(1 − *θN_m_*) × *f_i_* / (*h_f_* +Σ *f_i_*). The first term represents the number of mature eggs produced by the female, which decreases with the number of males that she rejected before the mating, *N_m_* (*N_m_* ≤ 1/*θ*), and *θ* is the mate search cost. The second term represents the egg fertility rate and the survival rate of newborn offspring, which increases with the paternal investment, *f_i_*, and saturates to 1. The parameter *h_f_* determines the speed of saturation. Then, surviving newborn individuals (i.e. juveniles) experience density-dependent mortality in which per capita survival probability is *K*/(*K*+*N*), where *N* is the number of juveniles in the population, and *K* is the population density at which survival rate becomes 1/2.

## Results

### Conditions for the existence of multiple alternative stable evolutionary equilibria

I first explored conditions for the evolution and maintenance of exaggerated male sexual displays and for the existence of multiple stable evolutionary equilibria. For this purpose, I conducted simulations of evolutionary dynamics in a closed population for 100,000 generations. The simulations start from an artificial initial population consisting of 100 individuals with the ancestral genome that contains no mutation, whose trait values are 0 for all traits.

Evolutionary dynamics in a closed population reached a stasis after one of male sexual displays became exaggerated to function as an honest signal of male quality (Fig. S2). In such an equilibrium state, an exaggerated male sexual display incurring fitness cost (i.e., the reduction of resource allocation to paternal investment, *f_i_*) was maintained by female mate preference causing sexual selection for the exaggerated sexual display. The female mate preference was maintained because there were direct fitness benefits of choosing males with an exaggerated sexual display. That is, males’ paternal investments to offspring fitness (*f_i_*) positively correlated with the exaggeration levels of their sexual displays (Σ*_s_*|*D_si_*|) because these male traits both increased with their resource budget, which was subject to environmental variance (Fig. S2c). Although the tradeoff in resource allocation contributes to generating a negative correlation between Σ*_s_*|*D_si_*| and *f_i_* among males with different resource allocation strategies (*d_si_*), this effect was not pronounced when genetic variation in *d_si_* was small compared to the environmental variance of resource budget, *_e_*.

Regardless of the preference function model, there were four possible combinations of male and female trait values to which evolution could converge (i.e. alternative stable evolutionary equilibria) (Figs. 1a, S5a, S6a). This was because exaggerated sexual displays could be generated by either overexpression or underexpression of either one of two display traits. The existence of multiple stable evolutionary equilibria in our model is consistent with a previous model considering direct benefits of mate choice [15]. Although evolutionary dynamics starting from the artificial initial state with the cryptic sexual display traits could reach any one of the alternative stable equilibria, mating displays and mate preference did not change for long times once a population reached one of stable equilibria (Fig. 1a, S5a, S6a). Therefore, simulation results suggest that allopatric speciation is unlikely to occur if the common ancestral species already has reached one of stable equilibria; natural and sexual selection stabilizing each equilibrium in turn hinder evolutionary divergence of mate preference and sexual displays between geographically isolated populations.

**Figure 1.**
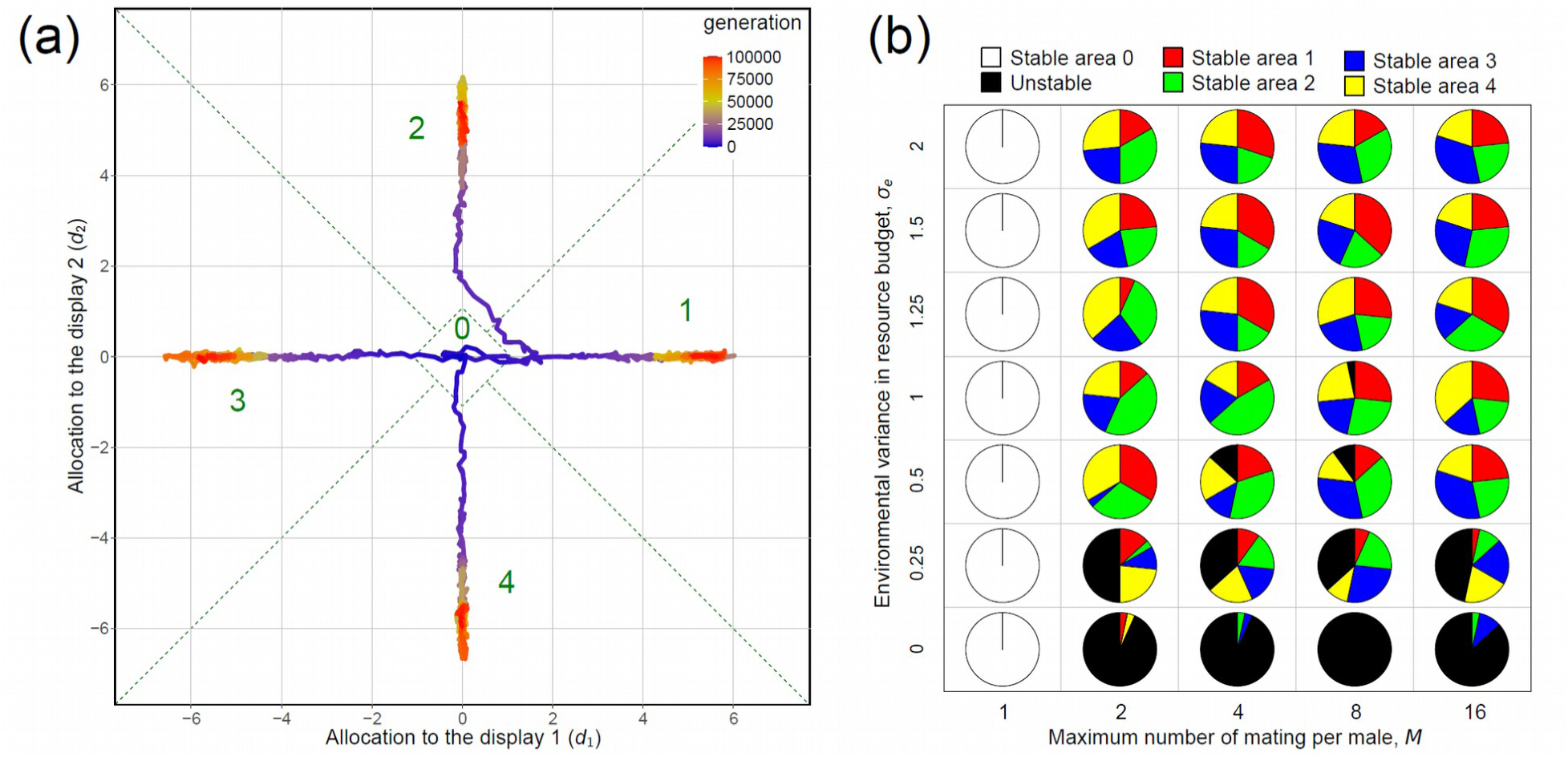
Alternative stable equilibria in evolutionary dynamics of two male sexual display traits. The simulations assume the beta preference function model. (a) Evolutionary trajectories of male genetic traits *d*_1_ and *d*_2_ in four simulations starting from the same artificially selected initial state (*d*_1_ = *d*_2_ = 0). Mean values of the traits across all males in the population, ⟨***d***⟩ = (⟨*d*_1_⟩, ⟨*d*_2_⟩), are shown. Colors indicate time (generation) in each simulation (blue: earlier; red: later). Green dotted lines show boundaries between five areas that are defined to categorize equilibria of evolutionary dynamics (see main text). Parameters were set to the default values (Table S1). (b) Results of simulations with systematically varied values of the environmental variance of resource budget, *_e_*, and the upper limit number of mating per male individual, *M* (other parameters were set to the default values). Pie chart for each set of parameter values shows frequencies that ⟨***d***⟩ in the population stayed in trait-areas 0, 1, 2, 3, and 4 for more than 100,000 generations (*n* = 30 simulation replications). Simulation results were categorized as “unstable” when ⟨***d***⟩ in the population did not stay in a single trait-area for more than 100,000 generations.

Alternative stable equilibria existed under conditions where sexual selection led to the evolution and maintenance of biased female mate preference and exaggerated male displays. Figure 1b summarizes results of simulations with systematically varied values of two parameters: the environmental variance in resource (budget (*_e_*) and the maximum number of mating per male individual (*M_m_*). For each parameter set, 30 simulation replications were conducted. Among them, four alternative stable equilibria were identified when *_e_* (was not small (*_e_* > 0.5) and *M_m_* was larger than 1. When *_e_* was small, the direct benefit of female mate choice was small because paternal investment did not vary greatly among males; hence, the female mate preference for exaggerated sexual displays was unlikely to evolve. When *M_m_* = 1, the mating system was monogamy and the disparity of mating success among males was small, which means that sexual selection on males was absent or very weak. These results were not qualitatively changed under different preference function models (Figs. S5, S6). The default values of parameters for simulations considering hybridization are selected from the parameter sets with which the evolution of exaggerated sexual display is possible; under such conditions, there are always four alternative stable evolutionary equilibria in this model.

### Simulation scenario 1: Hybridization in a secondary contact zone

Evolutionary dynamics caused by hybridization was simulated by considering a scenario where a common ancestral species is divided into two allopatric populations that subsequently come into secondary contact after a certain period of isolation. To eliminate the influence of the artificial initial state on simulation results, a burn-in period of 100,000 generations was provided at the beginning of simulation, during which the common ancestral species reached one of stable equilibria of evolutionary dynamics. Subsequently, the common ancestral population is divided into two geographically isolated populations. Independent evolution of two populations was simulated for *T*_0_ generations. Then, I simulated secondary contact in the third site (i.e. the contact zone) receiving recurrent immigration from both parental populations. Every generation, the number of immigrants from each parental population is given as a random number drawn from a Poisson distribution with intensity *m*. Immigration occurs at the juvenile stage before they experience the density-dependent mortality. Since density regulation occurs after migration, the effective immigration rate decreases as population density in the contact zone increases, although the expected number of immigrants is constant over time.

Hybridization could promote speciation by promoting evolutionary shifts between different stable equilibria (Figs.2, S11 - S13). Hybrid speciation occurred through two steps. The first step was the evolutionary loss of an exaggerated sexual display. In the parental populations, a costly exaggerated male display was maintained because most females shared similar mate preferences to cause a strong directional sexual selection for the exaggerated display. In the secondary contact zone, however, hybridization gave rise to large standing variation, including transgressive phenotypes, through segregation and recombination of chromosomes from two genetically distinct parental lineages (i.e. admixture variation) (Figs.2, S11). The increased variation of female mate preference in the hybrid population disturbed the alignment of mate preference among females. Hence, hybridization mostly reduced the gradient of the reproductive fitness landscape for males around the average sexual display phenotype; in other words, sexual selection was weakened (Figs.2b, S11b). This was true under all three models of preference function (Figs. S12 - S13). At the same time, the increased variation in male genetic traits that control resource allocations to sexual displays (*d*_1_ and *d*_2_) weakened the positive correlation between the degree of display exaggeration (Σ*_s_*|*D_si_*|) and paternal investment (*f_i_*). Therefore, admixture variation weakened both the sexual selection for a costly exaggerated male sexual display and the direct benefit of female mate choice, which used to stabilize the original equilibrium state in the parental populations. Consistent with these observations, hybridization was mostly followed by the rapid evolution of a cryptic male display with low fitness cost and a female mate preference against the original male display of parental lineages (Fig.2, S11; the period immediately after the onset of secondary contact); the phenotypic changes in male display and female preference then reduced the gene flow from parental populations to the hybrid population to some extent. The second step for speciation was the evolution of novel male displays and female mate preference. If ongoing gene flow from parental lineages was low, standing variation of female mate preference declined in the hybrid population, and then sexual selection drove the evolution toward one of alternative stable equilibria again. Speciation took place if the hybrid population evolved to a previously unoccupied stable equilibrium, because the hybrid population was reproductively isolated from both parental lineages owing to the divergence in mate preference and sexual displays (Figs.2, S10, S11). Simulations with the exponential and Gaussian preference function models also confirmed that hybridization can lead to speciation with the same mechanism (Figs. S12, S13).

**Figure 2.**
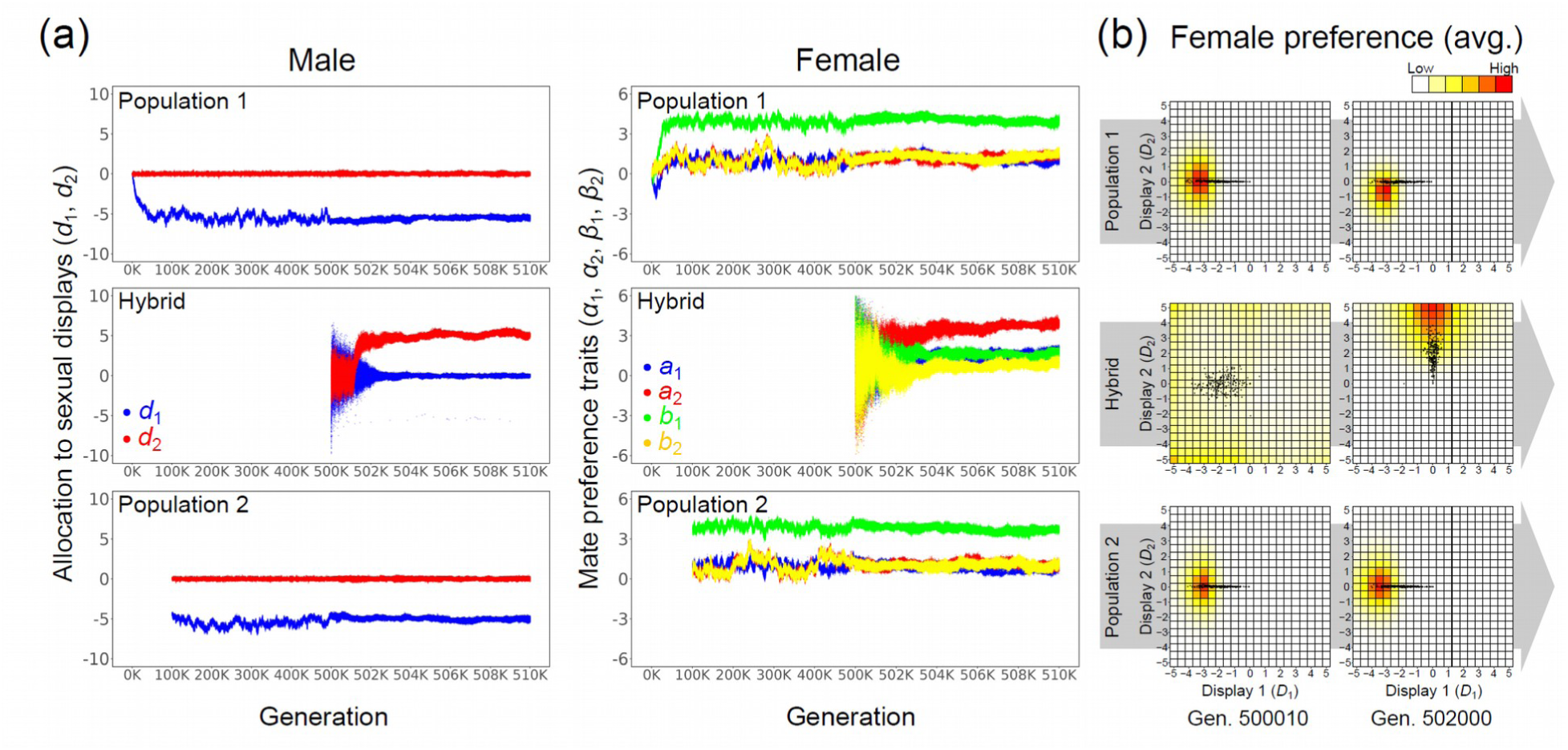
An example of simulation in which hybridization led to speciation. This simulation assumes the beta preference function model and the continuing immigration from parental-to the hybrid population. (a) Evolutionary dynamics of male genetic traits that determine relative resource allocation to two sexual displays (blue: *d*_1_, red: *d*_2_) and female genetic traits that determine the shape of mate preference function (blue: *a*_1_, red: *a*_2_, green: *b*_1_, yellow: *b*_2_). Trait values are shown for all male and female individuals in each population at each generation. The secondary contact period (generations 500K - 510K) is magnified along the horizontal axis. (b) Averaged female preference function across all female individuals (red colors) and sexual display phenotypes of all male individuals (dark gray points) are shown for each population at two time points (10 and 2000 generations after the secondary contact has begun at the generation 500,000). Horizontal and vertical axes show phenotypic values of two male sexual displays *D*_1_ and *D*_2_, and the coloration of each cell shows the probability that a randomly selected female will accept the male with the corresponding sexual display phenotypes upon a single encounter event. Parameters were set to the default values (Table S1).

Simulations with systematically varied values of the divergence time between parental lineages (*T*_0_) and the intensity of continuous immigration of parental lineages to hybrid population (*m*) revealed that hybridization can lead to several distinct outcomes depending on values of these parameters (Fig. 3a; see supplementary method for details of the categorization of evolutionary outcomes). First, hybridization did not lead to the evolutionary loss of the exaggerated male display in the first place if hybridization did not generate large enough standing variation for collapsing the original equilibrium state (Fig. S7). Accordingly, when the divergence time between hybridizing lineages was short (*T*_0_ ≤ 120,000 in Fig 3a), the hybrid population mostly maintained the same phenotypes as parental lineages (i.e., “no change”). Second, a cryptic sexual display could be maintained stably in the hybrid population after the evolutionary loss of the original exaggerated sexual display (Fig. S9). This outcome (“cryptic display”) occurred when intensive recurrent immigration from parental-to hybrid population (*m* > 8 in Fig. 3a) constantly generated high admixture variation in the hybrid population, which kept disturbing the sexual selection mechanisms for the evolution of exaggerated male displays. Similarly, intensive continuous immigration of parental lineages to the hybrid population (*m* > 8 in Fig. 3a) could drive continuous drift of male display phenotypes in the hybrid population (Fig. S8). In this case (“drift”), stabilization of any specific phenotype was inhibited due to the constant supply of genetic polymorphisms through recurrent hybridization. Third, if ongoing gene flow from parental lineages was moderate or lower (*m* ≤ 8 in Fig. 3a), the hybrid population evolved toward one of alternative stable equilibria again; if the hybrid population evolved to a previously unoccupied stable equilibrium, “speciation” or “incomplete speciation” occurred because the hybrid population was more or less reproductively isolated from both parental lineages owing to the divergence in mate preference and sexual displays (Figs.2, S10, S11).

**Figure 3.**
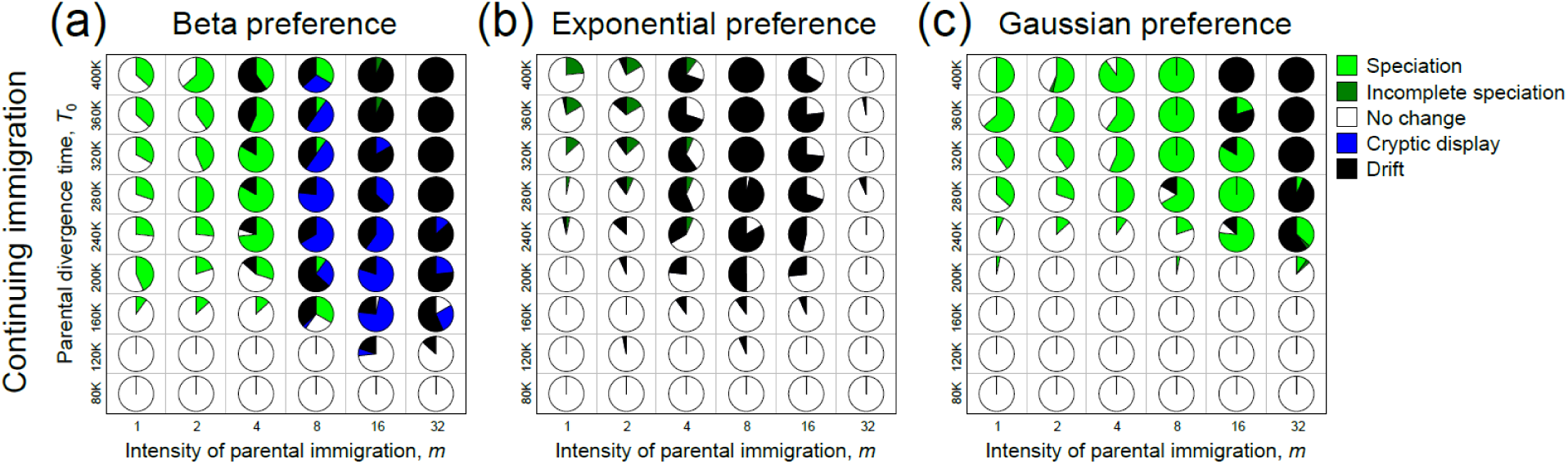
Conditions for hybrid speciation in simulations with various values of the divergence time between parental lineages (*T*_0_; vertical axis) and the intensity of immigration of parental lineages to the hybrid population (*m*; horizontal axis). The simulations assumed either the beta preference function model (a), the exponential preference function model (b), or the Gaussian preference function model (c). For each combination of parameter values, a pie chart shows frequencies of the following five categories of evolutionary outcomes in 30 simulation replicates: “speciation” (green); “incomplete speciation” (dark green); “no change” (white); “cryptic display” (blue); “drift” (black). Other parameters were set to the default values (Table S1).

Although hybridization promoted speciation with the same mechanism under all three preference function models, the conditions for hybrid speciation varied depending on the preference function model. In simulations with the Gaussian preference function model, both the overall likelihood of hybrid speciation and the conditions for hybrid speciation were similar to those with the beta preference function model; namely, hybrid speciation occurred fairly frequently with large *T*_0_ and moderate *m* (Fig. 3c). With the exponential preference function, on the other hand, the overall likelihood of hybrid speciation was reduced, and only the hybrid speciation with weak reproductive isolation (i.e., “incomplete speciation”) was observed. Moreover, hybrid speciation occurred only with low continuous parental immigration (*m* ≤ 4 in Fig. 3b) or without parental immigration after the establishment of the hybrid population (Fig. S14). The evolution of strong reproductive isolation did not occur with the exponential preference function model probably because the mate search cost inhibits the evolution of strongly biased mate preferences. Since the exponential preference function can express only open-ended preferences, strong preferences under this model inevitably lead to strong mate search costs. This effect probably has prevented the evolution of strongly biased mate preferences in simulations with the exponential preference function model.

### Simulation scenario 2: Hybridization by a cycle of isolation and reconnection of habitable areas

In addition to the above scenario, I simulated hybridization caused by a cycle of fission and fusion of populations due to environmental changes. This scenario considered a meta-population consisting of four local patches connected by migration of individuals and four periods with different migration rates among patches. Firstly, as in scenario 1, the ancestral species was prepared by simulating evolution in a single population for 100,000 generations, and then the ancestral species was introduced into one of the four patches. In the second period of 10,000 generations, the per-capita migration rate was set to 0.1, which caused the ancestral species to spread to four patches. Migration occurs at the juvenile stage, and each migrating individual moves from the patch of birth to a randomly selected one of the other patches. Then, an environmental change occurs to cause the third period, the isolation period, during which migration rate was set to 0. After the isolation period of 300,000 generations, another environmental change occurs to cause the fourth period, the secondary contact period, during which the migration rate was set to *κ*. Speciation and sexual radiation could occur if four local populations evolved to occupy two or more stable equilibria of evolutionary dynamics.

This scenario could lead to sexual radiation (Fig. S15). In all simulations with this scenario, the ancestral phenotypes were maintained in all patches throughout the isolation period of 300,000 generations. In the secondary contact period, however, admixture variation could cause each local population to deviate from the ancestral equilibrium. Then, each local population could evolve toward one of four alternative stable equilibria. In some instances of simulation, local populations reached different evolutionary equilibria and were reproductively isolated from each other owing to phenotypic differentiation in mate preference and sexual displays. This process could form a sexual radiation involving three or more species with diverse exaggerated sexual displays within 10,000 generations (Fig. S15).

Then, I explored conditions conducive to sexual radiation by conducting simulations with systematically varied values of *κ* (the migration rate in the secondary contact period). Since the effect of hybridization on evolutionary dynamics was qualitatively similar with all three mate preference function models under the first simulation scenario, simulations of this scenario were conducted only with the beta preference function model to reduce the computational cost. Simulations revealed that secondary contact with moderate migration rates (0.0008 ≤ *κ* ≤ 0.002) are most conducive to sexual radiation (Fig. 4). With too large *κ* (*κ* ≥ 0.006), genetic and phenotypic differentiation between four local populations was not maintained due to substantial gene flow between them. On the other hand, too small *κ* (*κ* ≤ 0.0002) limited the opportunity of hybridization for generating standing genetic variation that can drive speciation.

**Figure 4.**
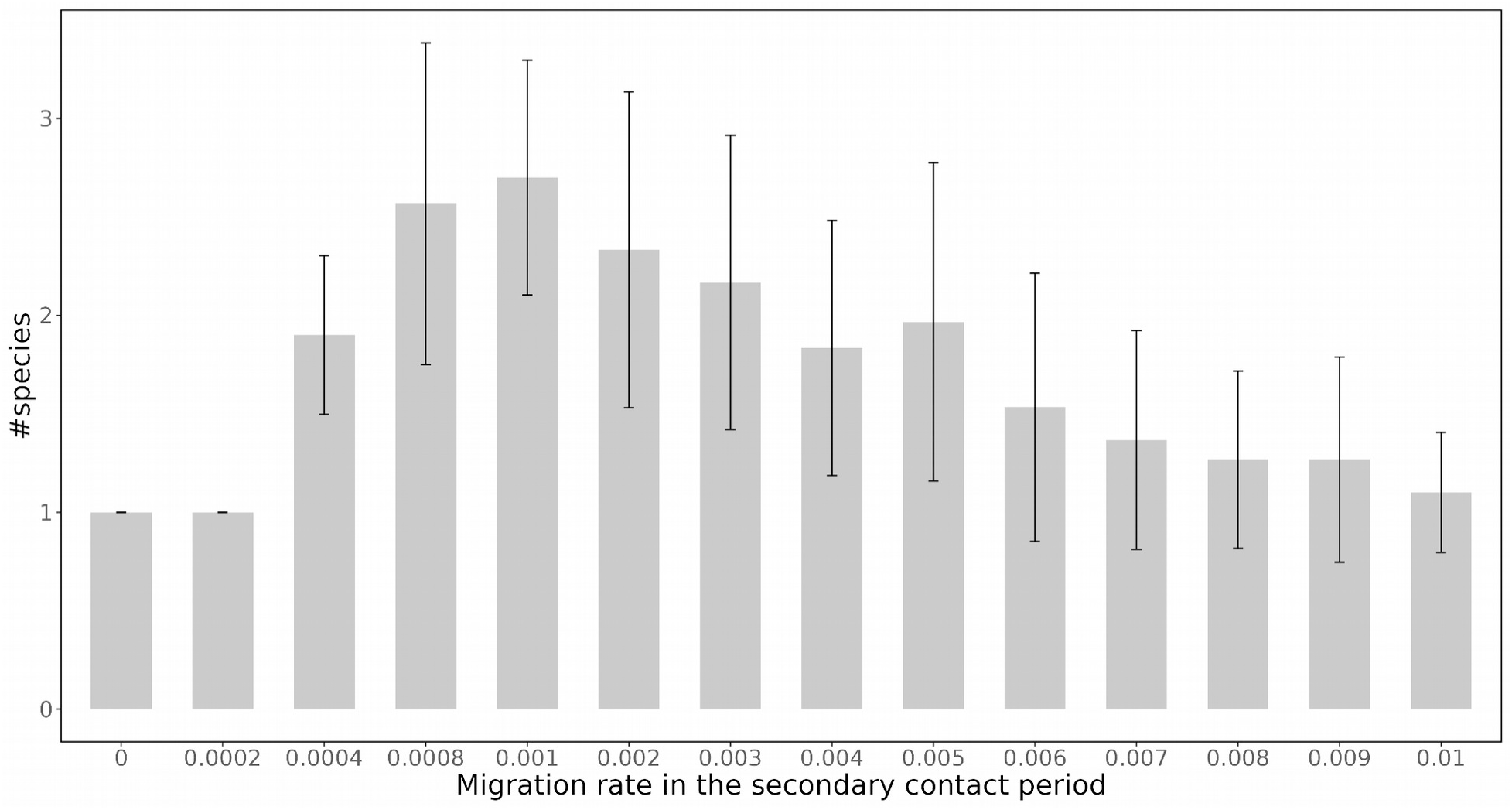
Conditions for sexual radiation in simulations with a cycle of isolation and reconnection of four habitable areas. Points show the number of species with different exaggerated sexual displays that have evolved in simulations with various values of the migration rate during the secondary contact period (horizontal). Bar plot shows the mean value across 30 simulation replicates with identical parameter settings and the error bar shows standard deviation. The method to count the number of species is described in the Supplementary Method. All other parameters were set to the default values (Table S1).

## Discussion

Sexual selection is considered an important driver of speciation and sexual radiation [1–3]. Supporting this view, existing theories of sexual selection suggest that evolutionary dynamics with sexual selection often have multiple alternative stable equilibria, since females can evolve preference for arbitrary male traits [15–17]. However, it remains unclear whether the presence of such alternative stable equilibria can explain speciation and sexual radiation, because little is known about how a group of species occupying different stable equilibria can be formed in the first place. Evolutionary simulations in the present study demonstrate that hybridization can bridge the presence of multiple stable equilibria and the incipient formation of sexual radiation. That is, hybridization can trigger the formation of new species that occupy previously unoccupied evolutionary equilibria that sexual selection generates.

Admixture variation of female mate preference and male sexual display including transgressive segregants played key roles in the rapid evolution of novel mate preference and sexual displays in the hybrid speciation process. Transgressive segregation could occur in the simulations because genomes of two parental lineages had independently accumulated mutations with compensating effects, while keeping their phenotypes constant. In the hybrid population, pairs of mutations with compensating phenotypic effects that have fixed within each lineage could segregate from each other to form novel genotypes that aggregate alleles with similar phenotypic effects inherited from different parental lineages. This process could create transgressive segregants with novel extreme phenotypes, despite the fact that parental lineages were phenotypically identical. With all three preference function models considered in this study, transgressive segregation of female mate preference tended to weaken the sexual selection for the exaggerated male sexual display of the parental lineage, thereby breaking down the balance between sexual selection for and natural selection against the costly exaggerated sexual display. This effect promoted the evolution of less exaggerated sexual displays in early hybrid populations, which opened up an opportunity for the subsequent evolution toward previously unoccupied stable equilibria.

These results extend a previous theory positing that transgressive segregation of mating traits (e.g., mating season, mating habitat selection, genitalia shapes, sexual display, and mate preference) can promote the evolution of reproductive isolation between the hybrid population and both parental lineages [22]. Although evolutionary simulations in this previous study provided clues to the incipient process of sexual radiation, the study was insufficient to explain sexual radiation for three reasons. First, their model did not intend to simulate the evolution of exaggerated sexual displays; they assumed no fitness benefits of mate choice that promote the evolution and maintenance of female mate preference for costly exaggerated sexual displays. Accordingly, only slightly costly sexual displays could evolve in their simulations. Second, there were only two alternative stable evolutionary equilibria under their model setting, with which the formation of more than two species was unfeasible. Third, their model did not consider evolution in multiple sexual display traits, let alone the formation of multiple species with qualitatively different sexual displays. For these reasons, the previous study did not explain the evolution of multiple species with diverse exaggerated sexual displays. The present study extends the previous study by considering multi-dimensional sexual displays and a fitness benefit of mate choice, with which sexual selection alone causes evolutionary dynamics with more than two stable equilibria with distinct costly exaggerated sexual displays. Additionally, the present study considered two alternative geographic settings of hybrid population (secondary contact between two lineages and a cycle of fission and fusion of four populations) and three alternative models of mate preference (Gaussian-, exponential-, and beta preference function models), whereas the previous study considered only the secondary contact of two lineages and the Gaussian preference function. With these model extensions, results in the present study demonstrate not only the robustness of the arguments in the previous study [22] but also that an interplay between hybridization and sexual selection can promote rapid sexual radiation in which three or more species with distinct exaggerated sexual displays are formed.

The present study invoked a direct benefit of mate choice but did not explicitly consider indirect genetic benefits (e.g. good gene benefit), although the model inevitably involved the sexy son effect (or Fisherian selection), which occurs whenever both sexual displays and mate preferences are heritable. I did not focus on indirect genetic benefits of mate choice because they may not be sufficient for long-term maintenance of costly sexual displays. One reason for this is that indirect genetic benefits vanish when the evolution of mate preference leads to strong sexual selection that eliminates genetic variation among males to be selected by females (i.e., the lek paradox [48]). Since it remains unclear under what conditions indirect genetic benefits alone can maintain female mate preference for long term, interpretation of simulation results becomes difficult when indirect genetic benefit is the main driver of the evolution of mate preference. The lek paradox, however, does not matter when there is environmental (non-genetic) variance of male condition (e.g., the level of parasite load) affecting direct benefits of female mate choice (e.g., the risk of parasite infection through mating) [49]. The present study thus considered such a situation to improve the interpretability of simulation results. The condition for hybrid speciation driven by sexy-son and good-gene benefits of mate choice is an important open question to be investigated. Nonetheless, it would be reasonable to expect that transgressive segregation of female mate preference would promote hybrid speciation also when indirect genetic benefits of mate choice are the main drivers of the evolution of female mate preference. This is because the maintenance of a certain costly male sexual display would generally require the concordance of mate preference across females within the population. Hence, hybridization generating standing variation of female mate preference may generally hinder the maintenance of the original male sexual display, which in turn creates opportunities for the evolution of novel exaggerated sexual displays.

The probability of hybrid speciation depended on the preference function model. Namely, hybrid speciation was less likely to occur with the exponential preference function model than with the Gaussian and beta preference function models. This result is reasonable considering that the exponential preference function model can express only one-sided open-ended preferences, which means that females with any possible phenotypes of mate preference prefer exaggerated sexual displays (except for *p*_1_ = *p*_2_ = 0 causing random mating). Consequently, no matter how large the admixture variation of mate preference, sexual selection against the cryptic phenotypes of sexual display must persist in the hybrid population, suppressing the first step for hybrid speciation: the evolutionary loss of the original exaggerated male display of parental lineages. Gaussian and beta preference function models are not subject to this evolutionary constraint inhibiting hybrid speciation. These results indicate that the evolutionary dynamics after hybridization strongly depends on the relationship between genotypes and shapes of the preference function, but unfortunately, virtually nothing is known about this relationship even in model organisms of sexual selection to my knowledge.

Whether or not sexual radiation can occur should depend on the number of alternative stable evolutionary equilibria, which should increase with the number of potential traits that can evolve to become sexual displays. Additionally, the nature of the tradeoff limiting the expression of multiple sexual displays may be crucial for the stability of certain sexual displays. The present study adopted a natural assumption that each male possesses a limited resource budget and that increasing the investment to one of two sexual display traits automatically decreases the relative investment to another sexual display trait. Owing to this energetic trade-off, increasing the total investment to sexual displays decreases the absolute phenotypic effect of single alleles that additively affect the relative resource allocations to display traits. Accordingly, evolution of an exaggerated phenotype in one of two sexual display traits reduces the absolute phenotypic effects of mutations on another sexual display trait, thereby suppressing the evolution of the second exaggerated male display. Without this trade-off, evolutionary dynamics in two sexual display traits may not stop at certain equilibrium points, because any combinations of two sexual display phenotypes with the same level of resource demand will become neutrally stable equilibria. Under such a condition, relative weights for the two display traits in female mate preference will easily change due to genetic drift, causing continuing evolution of mate preference and sexual displays. This situation is outside the scope of our hypothesis.

The present study suggests that an interplay of hybridization and sexual selection can promote speciation if hybridization generates high enough standing variation in the mate preference function to modify the regime of intersexual selection. In line with hybridization promoting speciation by sexual selection, genomic evidence of past hybridization events is reported from many evolutionary radiations with diversification of sexual displays [12,13,29–36,50]. However, empirical data on how hybridization may affect mate preference phenotypes is scarce; genetic basis of mate preference function is largely unknown even in model systems of sexual selection and evolutionary radiations, and, to my knowledge, no previous study has tested whether hybridization can lead to transgressive segregation of mate preference function. I propose that an important future direction is to empirically test whether hybridization can generate various novel shapes of the mate preference function.

## Conclusion

Previous speciation theories suggest that sexual selection alone is not likely to promote speciation-with-gene-flow [51], although sexual selection can spur the ecological speciation process (e.g. sensory drive speciation) [52,53]. Hence, sexual radiations without ecological diversification seem to contradict with the theoretical expectation. The present study proposes a plausible resolution for this conundrum: sexual selection can drive non-ecological speciation in parapatry when hybridization catalyzes the evolutionary shifts between alternative stable equilibria of evolution that sexual selection generates. This speciation mechanism can also offer an alternative route to adaptive radiation: sexual radiation followed by ecological character displacement between species. Thus, we may need to reevaluate the power of sexual selection as a primary driver of evolutionary radiation.

## Acknowledgments

Simulations in this study were carried out with a super computer of Institute of Statistical Mathematics (ISM) under the ISM Cooperative Research Program (2023-ISMCRP-0010). I thank members of the Fish Ecology & Evolution department of EAWAG, Kawata laboratory of the Graduate School of Life Sciences of the Tohoku University, and Kitano laboratory of the National Institute of Genetics for fruitful discussions.

## Funding

This study was supported by the JSPS Overseas Research Fellowship, the JSPS Research Fellowships for Young Scientists (20J00583), and Japan Society for the Promotion of Science KAKENHI grants (22K15163).

## Supplementary table

**Table S1.** Model Parameters.

| Definition | Symbol | Default | Alternative values examined |
| --- | --- | --- | --- |
| The number of loci that can affect each genetic trait | $L$ | 500 | - |
| Variance of mutation effect | $\sigma_m$ | 0.05 | - |
| Mate search cost | $\theta$ | 0.001 | - |
| The level of paternal investment at which the egg survival rate reaches 1/2 | $h_f$ | 0.25 | - |
| The maximum size of resource budget | $E_{max}$ | 5 | - |
| Environmental variance of male resource budget | $\square_e$ | 1.0 | 0.0, 0.25, 0.5, 1.25, 1.5, 2.0 |
| Upper limit number of mating per male individual | $M$ | 2 | 1, 2, 4, 8, 16 |
| The population density at which the juvenile survival rate becomes 1/2 | $K$ | 500 | - |
| The duration of allopatric evolution of parental lineages (generations) | $T_0$ | 300,000 | 80,000, 120,000, 160,000, 200,000, 240,000, 280,000, 320,000, 360,000, 400,000 |
| Initial ratio of two parental lineages (one-time immigration) | $r_0$ | 0.5 | 0.1, 0.2, 0.3, 0.4 |
| Expected number of immigrants per generation (continuing immigration) | $m$ | 0 | 1, 2, 4, 8, 16, 32 |
| Per capita migration rate in the secondary contact period (a cycle of isolation & reconnection) | $\kappa$ | 0 | 0.0002, 0.0004, 0.0008, 0.001, 0.002, 0.003, 0.004, 0.005, 0.006, 0.007, 0.008, 0.009, 0.01 |

## Supplementary figures

**Figure S1.**
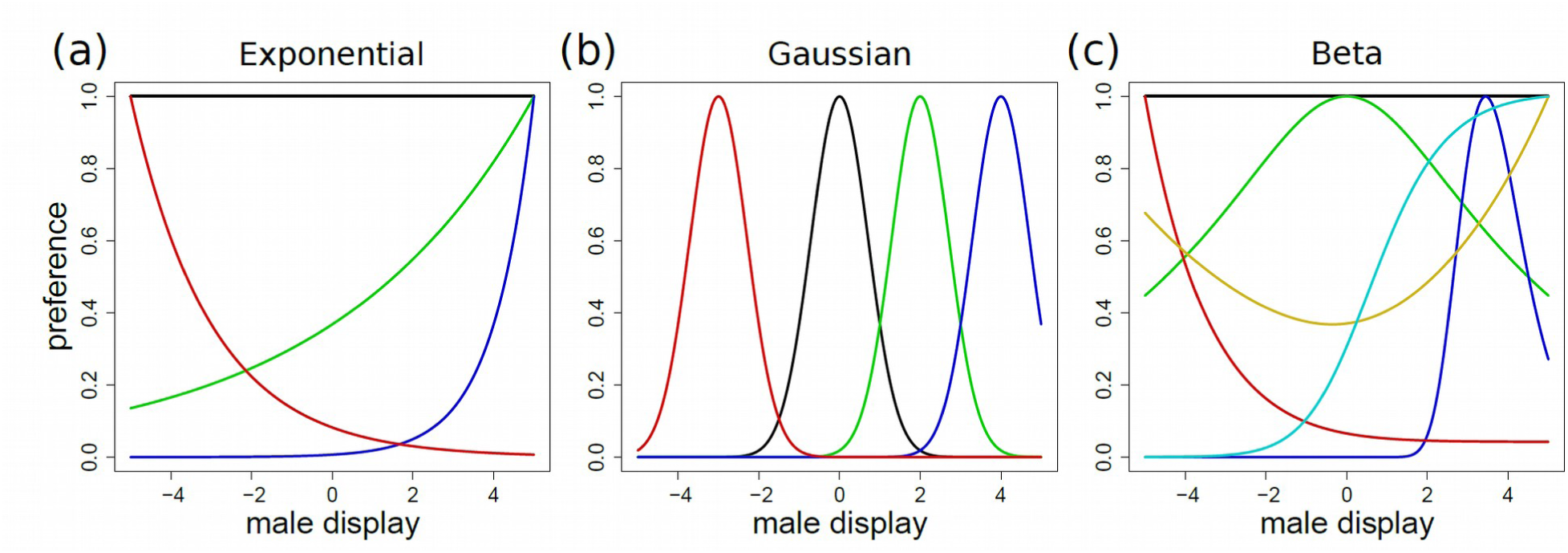
Three preference function models considered in this study. (a) The exponential preference function. Four curves show example shapes of the preference function with different values of the female trait, *p* (black: *p* = 0; green: *p* = 0.2; blue: *p* = 4; red: *p* = −0.5). (b) The Gaussian preference function (black: *x* = 0; green: *x* = 2; blue: *x* = 4; red: *x* = −3). (c) The beta preference function (black: *a* = 0, *b* = 0; green: *a* = 0.2, *b* = 0.2; blue: *a* = 4, *b* = 1; red: *a* = −1, *b* = 0; yellow: *a* = −0.2, *b* = −0.3; cyan: *a* = 1, *b* = 0).

**Figure S2.**
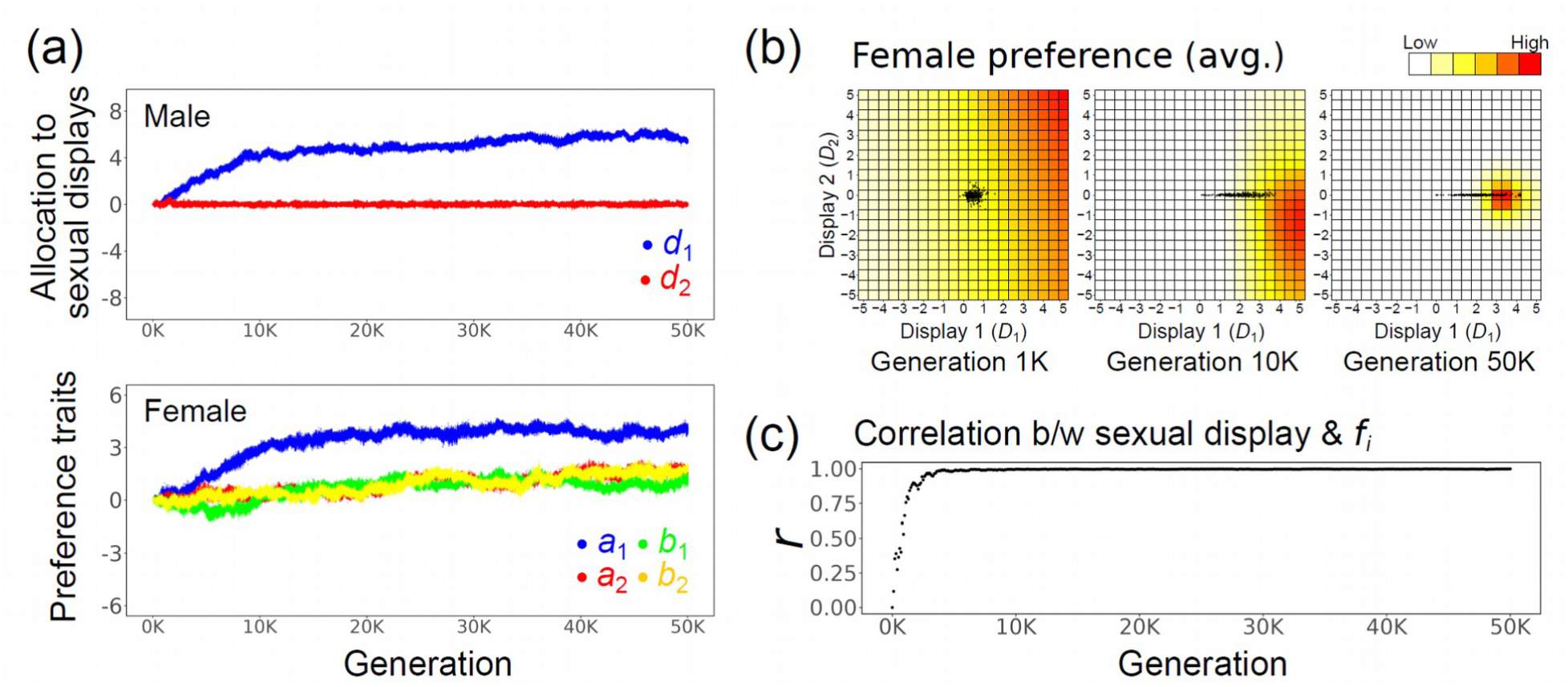
An example of evolutionary dynamics in a closed population. The simulation assumes the beta preference function model. (a) Evolutionary dynamics of male genetic traits that determine relative resource allocation to two sexual displays (blue: *d*_1_, red: *d*_2_) and female genetic traits that determine the shape of mate preference function (blue: *a*_1_, red: *a*_2_, green: *b*_1_, yellow: *b*_2_). Trait values are shown for all male and female individuals at each generation. (b) Averaged female preference function across all female individuals (red colors) and sexual display phenotypes of all male individuals (dark gray points) are shown for three time points (generations 1000, 10,000, and 50,000). Horizontal and vertical axes show phenotypic values of two male sexual displays *D*_1_ and *D*_2_. Color of cells show average female preference (i.e., the probability of acceptance upon an encounter) for each combination of male display values relative to that for the most preferred male display (red). (c) Correlation coefficient between the magnitude of display exaggeration (Σ*_j_*|*D_ji_*|) and the level of paternal investment on offspring fitness (*f_i_*) across all male individuals at each generation. Parameters were set to the default values (Table S1). Simulation examples with the exponential and Gaussian preference function models are in Figs. S3 and S4.

**Figure S3.**
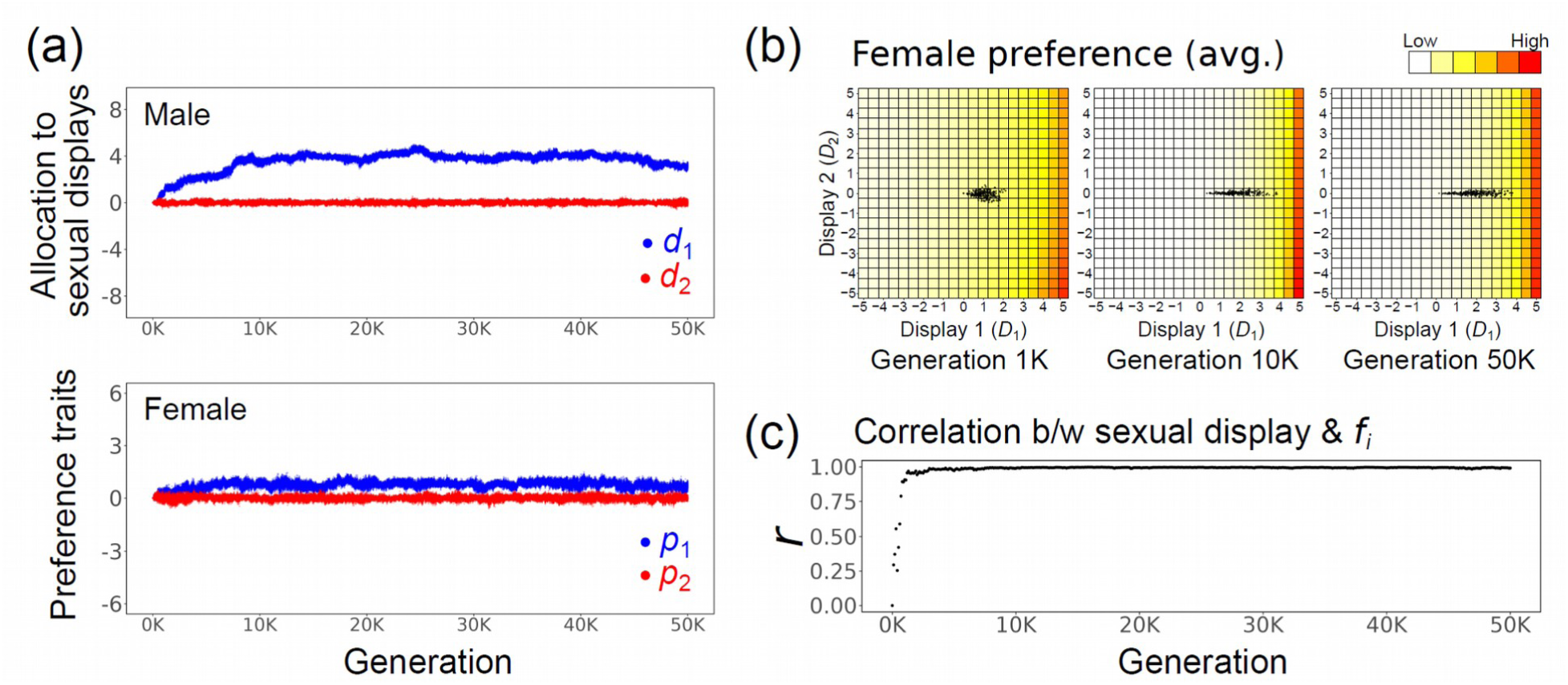
An example of hybrid speciation in a simulation assuming the exponential preference function model and continuing immigration from parental-to the hybrid population. Evolutionary dynamics is shown in the same format as Fig. S2, except that there are only two female traits (blue: *p*_1_, red: *p*_2_) in this simulation. Parameters were set to the default values (Table S1).

**Figure S4.**
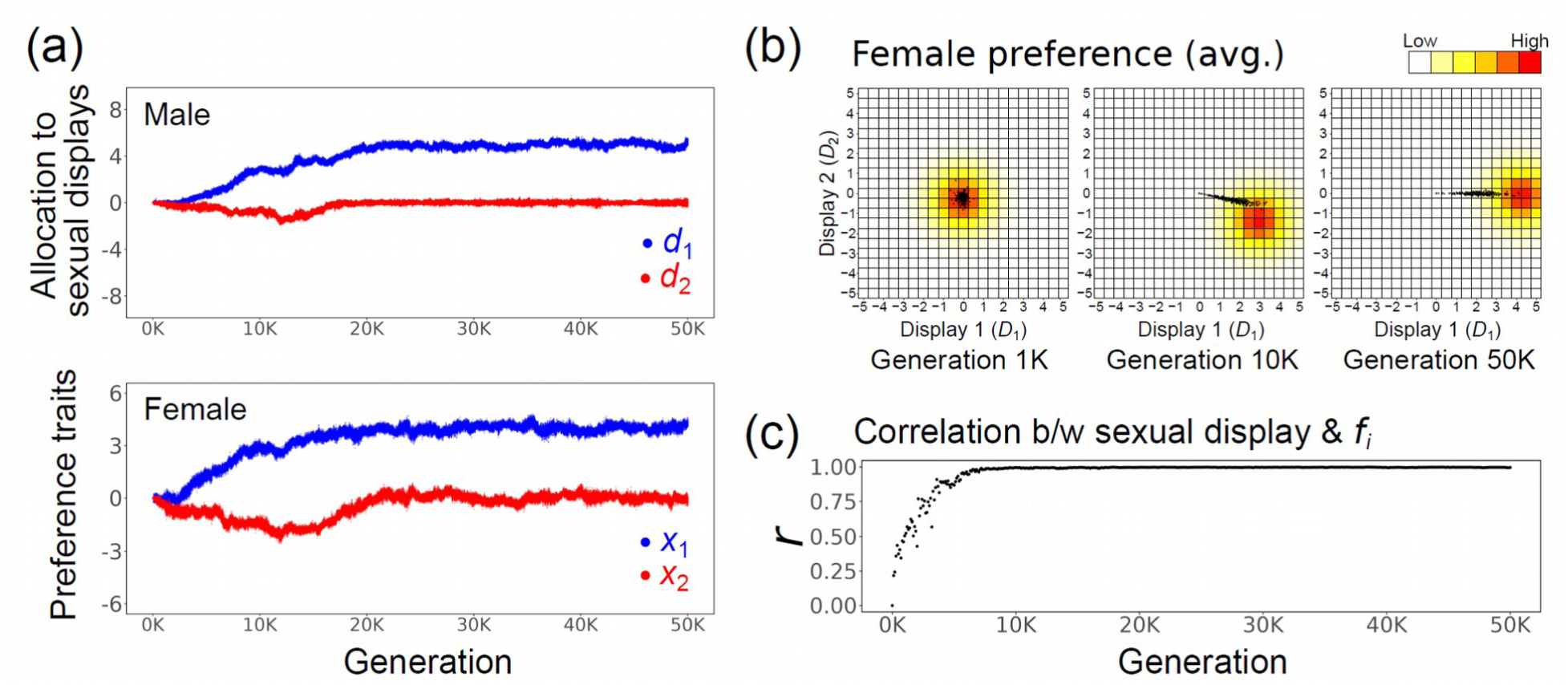
An example of hybrid speciation in a simulation assuming the Gaussian preference function model and continuing immigration from parental-to the hybrid population. Evolutionary dynamics is shown in the same format as Fig. S2, except that there are only two female traits (blue: *x*_1_, red: *x*_2_) in this simulation. Parameters were set to the default values (Table S1).

**Figure S5.**
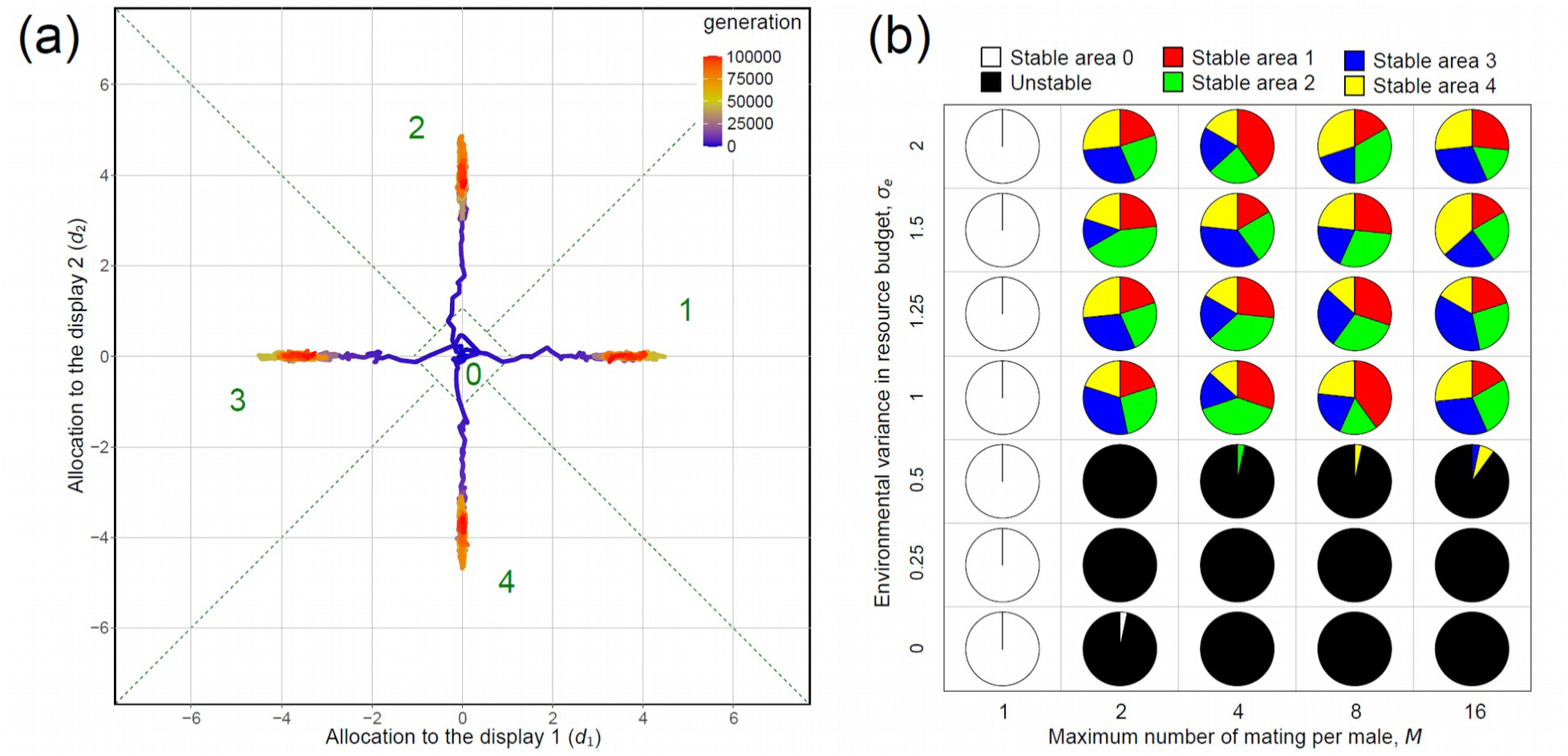
Alternative stable equilibria in evolutionary dynamics of two male sexual display traits in simulations assuming the exponential preference function model. (a) Evolutionary trajectories of male genetic traits *d*_1_ and *d*_2_ in four simulations starting from the same artificially selected initial state (*d*_1_ = *d*_2_ = 0) are shown in the same format as Figure 1a. (b) Results of simulations with systematically varied values of the environmental variance of resource budget, *_e_*, and the upper limit number of mating per male individual, *M* are shown in the same format as Figure 1b (other parameters were set to the default values).

**Figure S6.**
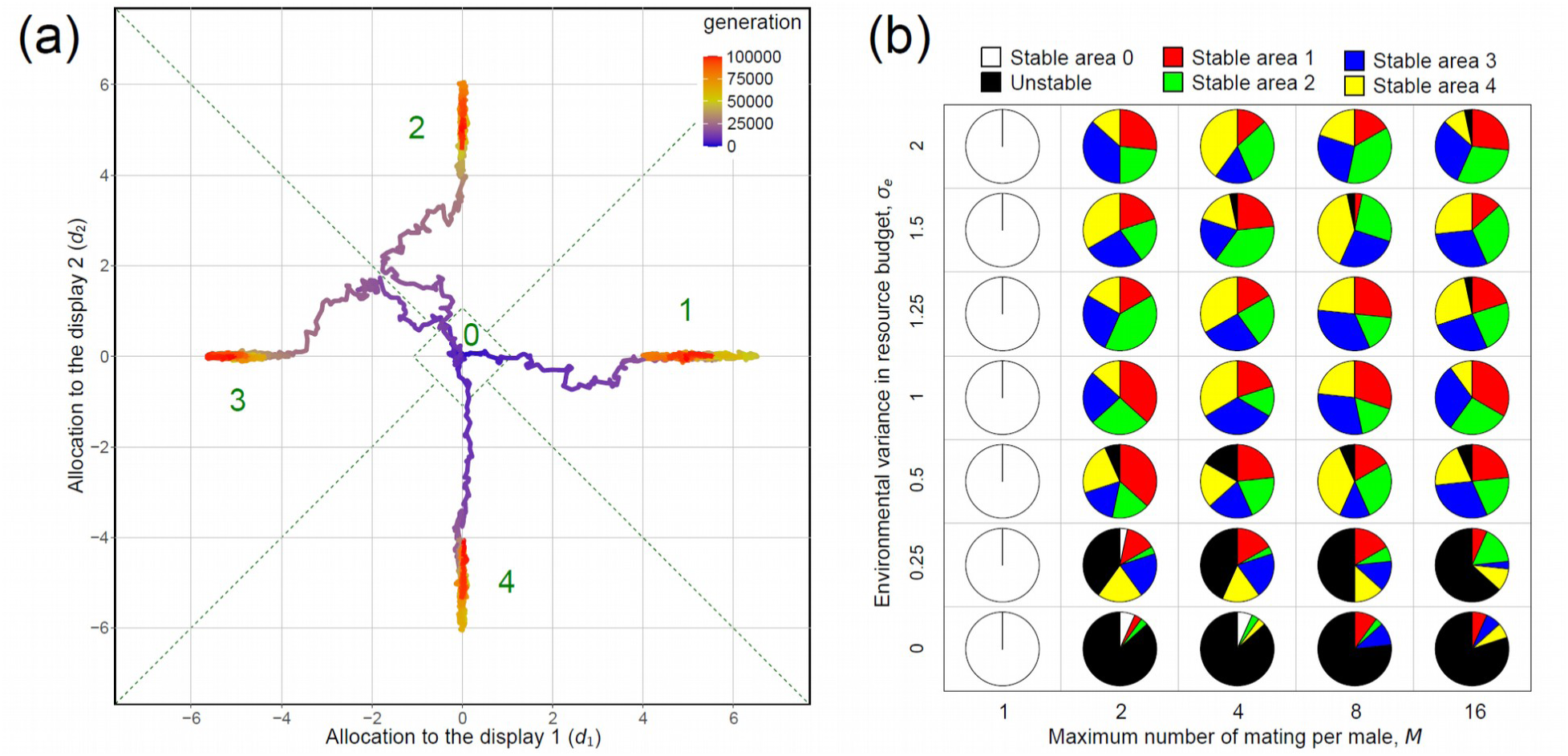
Alternative stable equilibria in evolutionary dynamics of two male sexual display traits in simulations assuming the Gaussian preference function model. (a) Evolutionary trajectories of male genetic traits *d*_1_ and *d*_2_ in four simulations starting from the same artificially selected initial state (*d*_1_ = *d*_2_ = 0) are shown in the same format as Figure 1a. (b) Results of simulations with systematically varied values of the environmental variance of resource budget, *_e_*, and the upper limit number of mating per male individual, *M* are shown in the same format as Figure 1b (other parameters were set to the default values).

**Figure S7.**
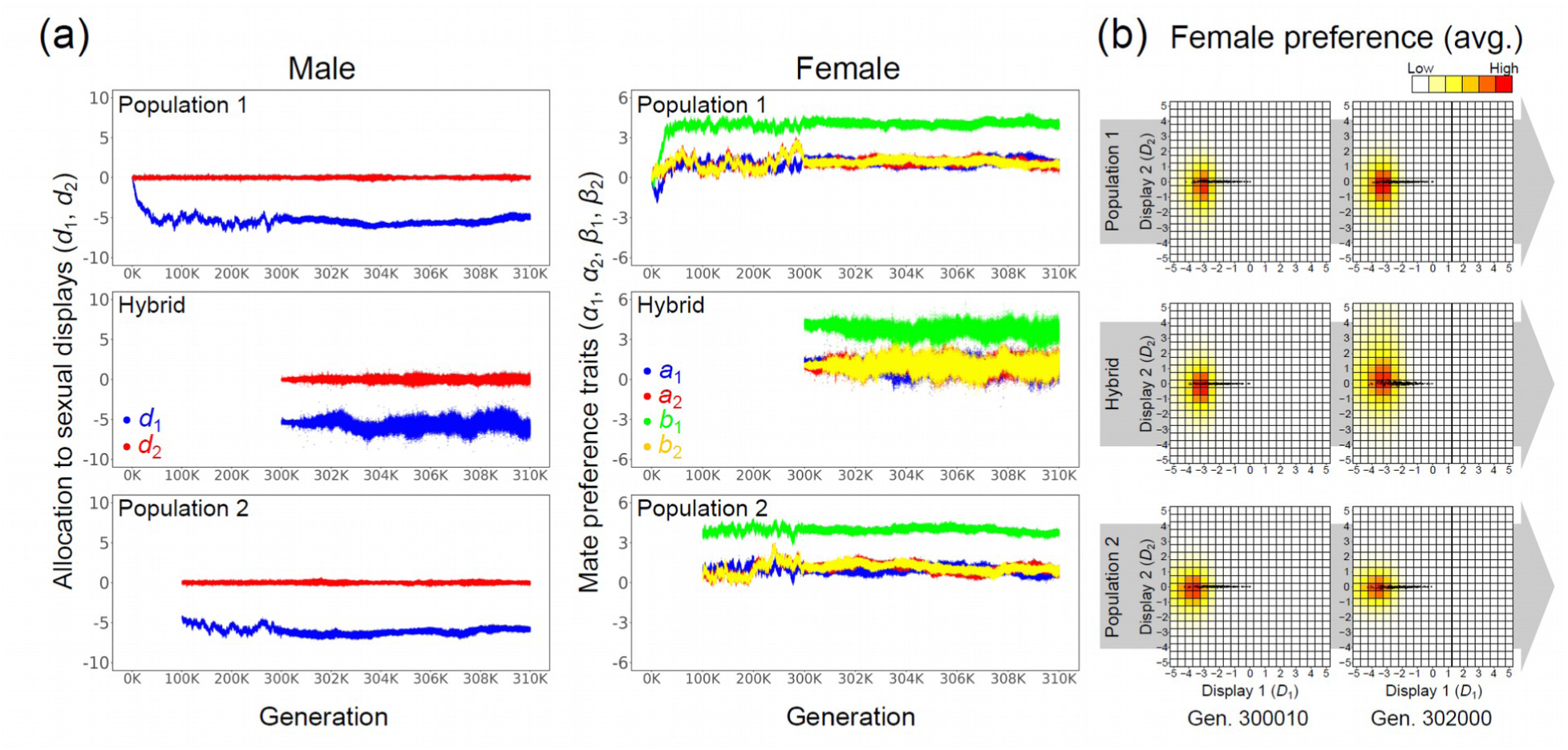
An example of simulation in which hybridization did not cause significant evolutionary change in male sexual displays (“no change”). The simulation assumed the beta preference function model and the continuing immigration from parental-to the hybrid population. (a) Evolutionary dynamics of two male genetic traits (blue: *d*_1_, red: *d*_2_) and four female genetic traits (blue: *a*_1_, red: *a*_2_, green: *b*_1_, yellow: *b*_2_) are shown in the same format as Figure 2a. (b) Averaged female preference function across all female individuals (red colors) and sexual display phenotypes of all male individuals (dark gray points) are shown in the same format as Figure 2b. Parameters were set to the default values (Table S1).

**Figure S8.**
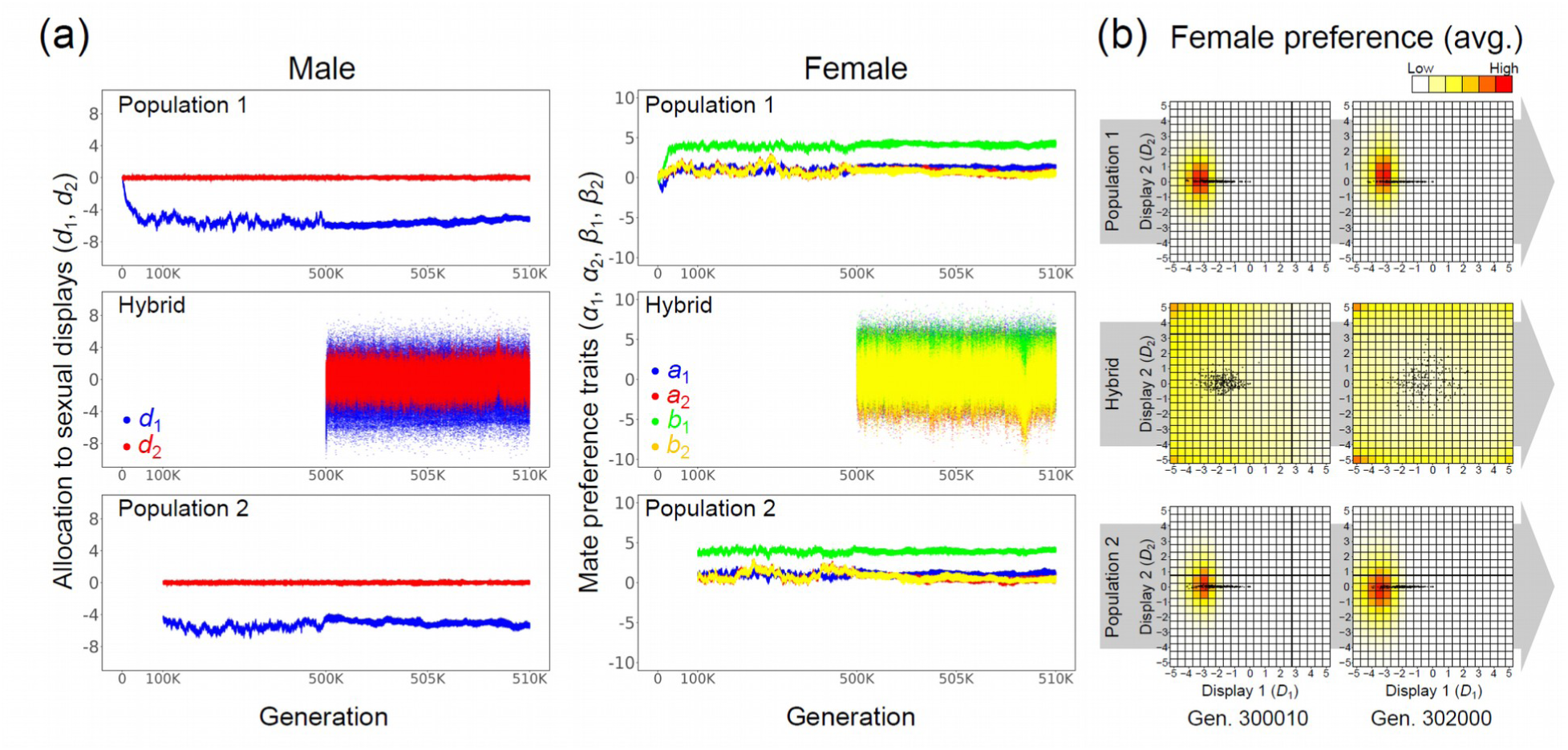
An example of simulation in which male traits in the hybrid population continued to drift until the end of simulation (“drift”). The simulation assumed the beta preference function model and the continuing immigration from parental-to the hybrid population. (a) Evolutionary dynamics of two male genetic traits (blue: *d*_1_, red: *d*_2_) and four female genetic traits (blue: *a*_1_, red: *a*_2_, green: *b*_1_, yellow: *b*_2_) are shown in the same format as Figure 2a. (b) Averaged female preference function across all female individuals (red colors) and sexual display phenotypes of all male individuals (dark gray points) are shown in the same format as Figure 2b. Parameters were set to the default values (Table S1).

**Figure S9.**
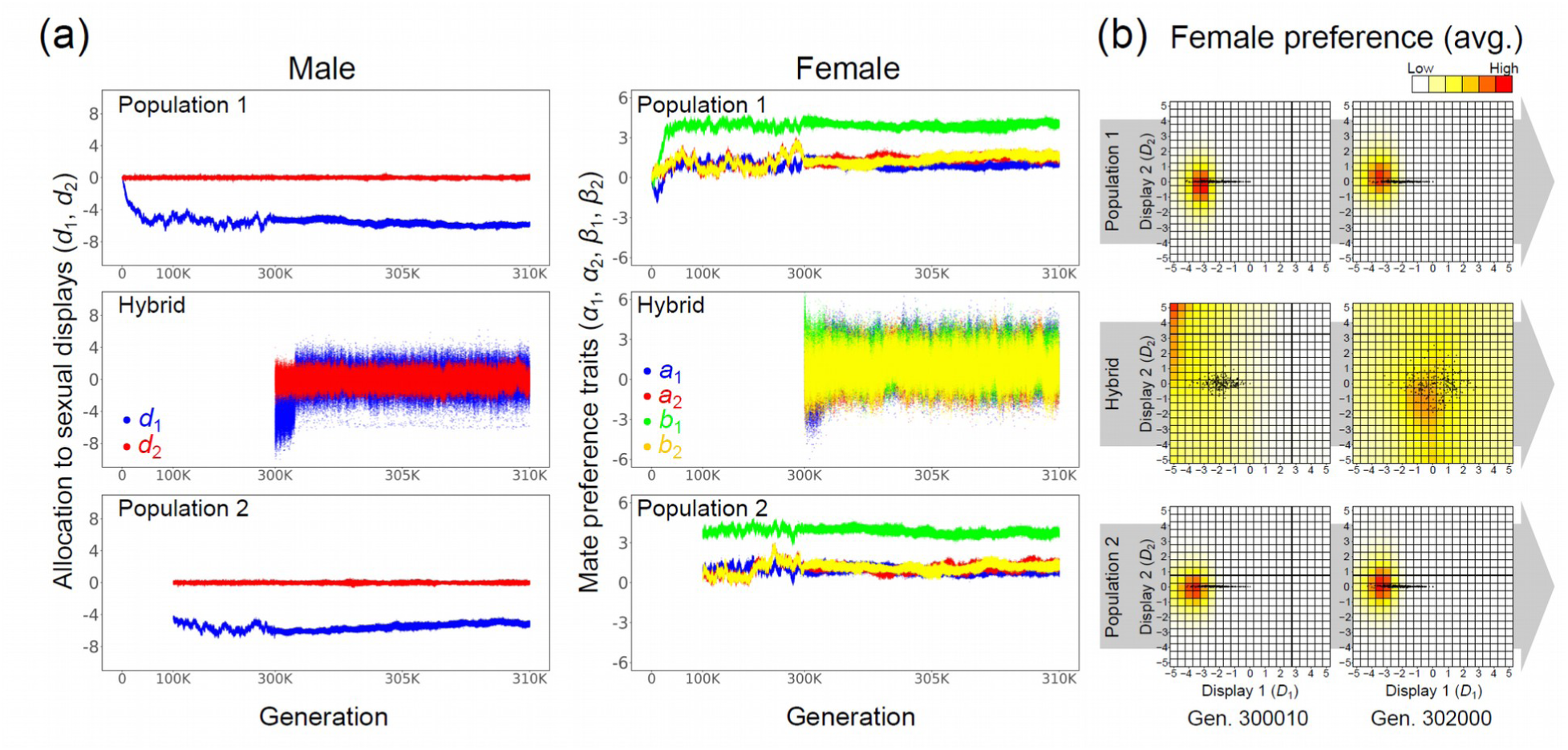
An example of simulation in which hybridization led to evolution and maintenance of the cryptic sexual display (“cryptic display”). The simulation assumed the beta preference function model and the continuing immigration from parental-to the hybrid population. (a) Evolutionary dynamics of two male genetic traits (blue: *d*_1_, red: *d*_2_) and four female genetic traits (blue: *a*_1_, red: *a*_2_, green: *b*_1_, yellow: *b*_2_) are shown in the same format as Figure 2a. (b) Averaged female preference function across all female individuals (red colors) and sexual display phenotypes of all male individuals (dark gray points) are shown in the same format as Figure 2b. Parameters were set to the default values (Table S1).

**Figure S10.**
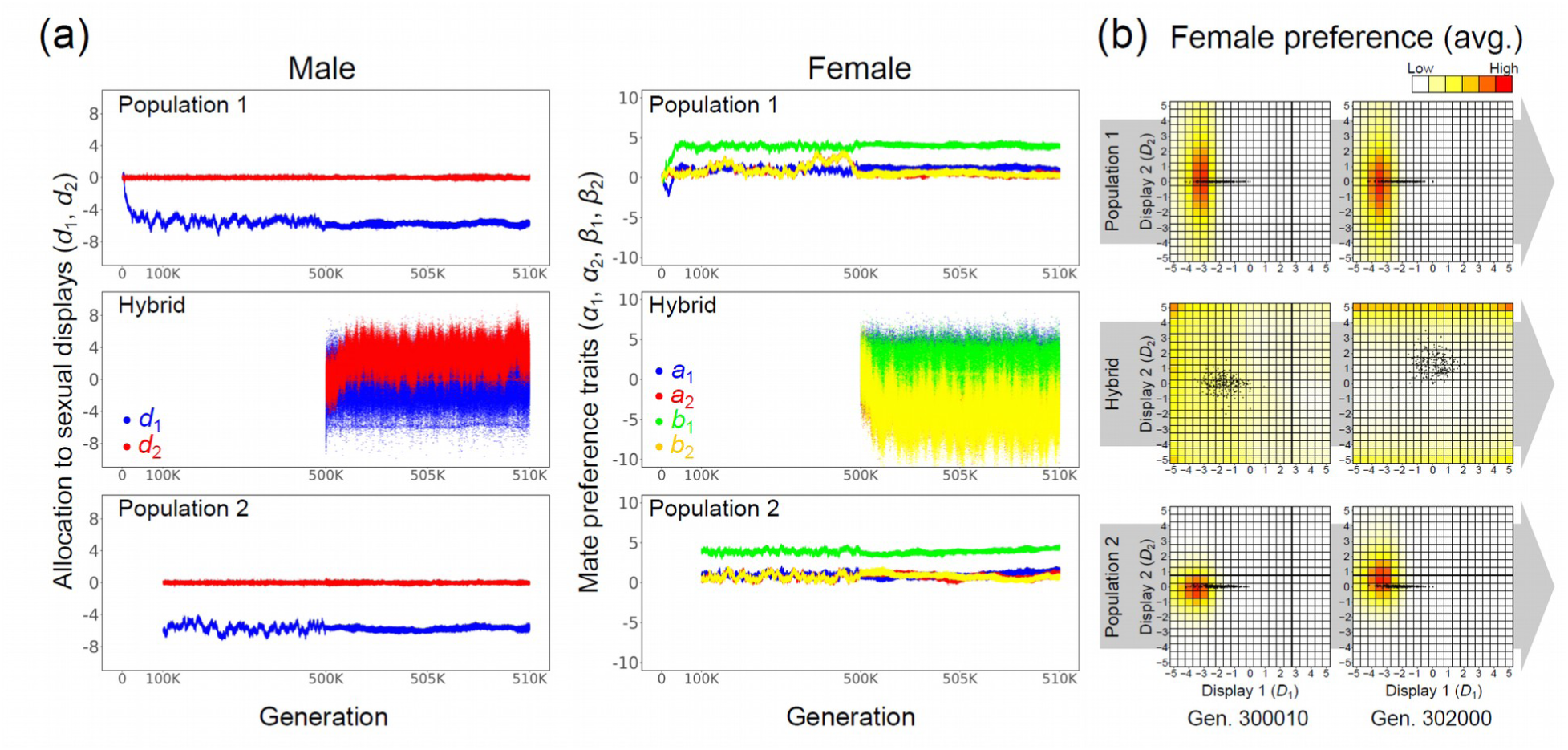
An example of simulation in which the hybrid population evolved novel phenotypes of male sexual displays and female mate preference but was not strongly reproductively isolated from the parental lineages (“incomplete speciation”). The simulation assumed the beta preference function model and the continuing immigration from parental-to the hybrid population. (a) Evolutionary dynamics of two male genetic traits (blue: *d*_1_, red: *d*_2_) and four female genetic traits (blue: *a*_1_, red: *a*_2_, green: *b*_1_, yellow: *b*_2_) are shown in the same format as Figure 2a. (b) Averaged female preference function across all female individuals (red colors) and sexual display phenotypes of all male individuals (dark gray points) are shown in the same format as Figure 2b. Parameters were set to the default values (Table S1).

**Figure S11.**
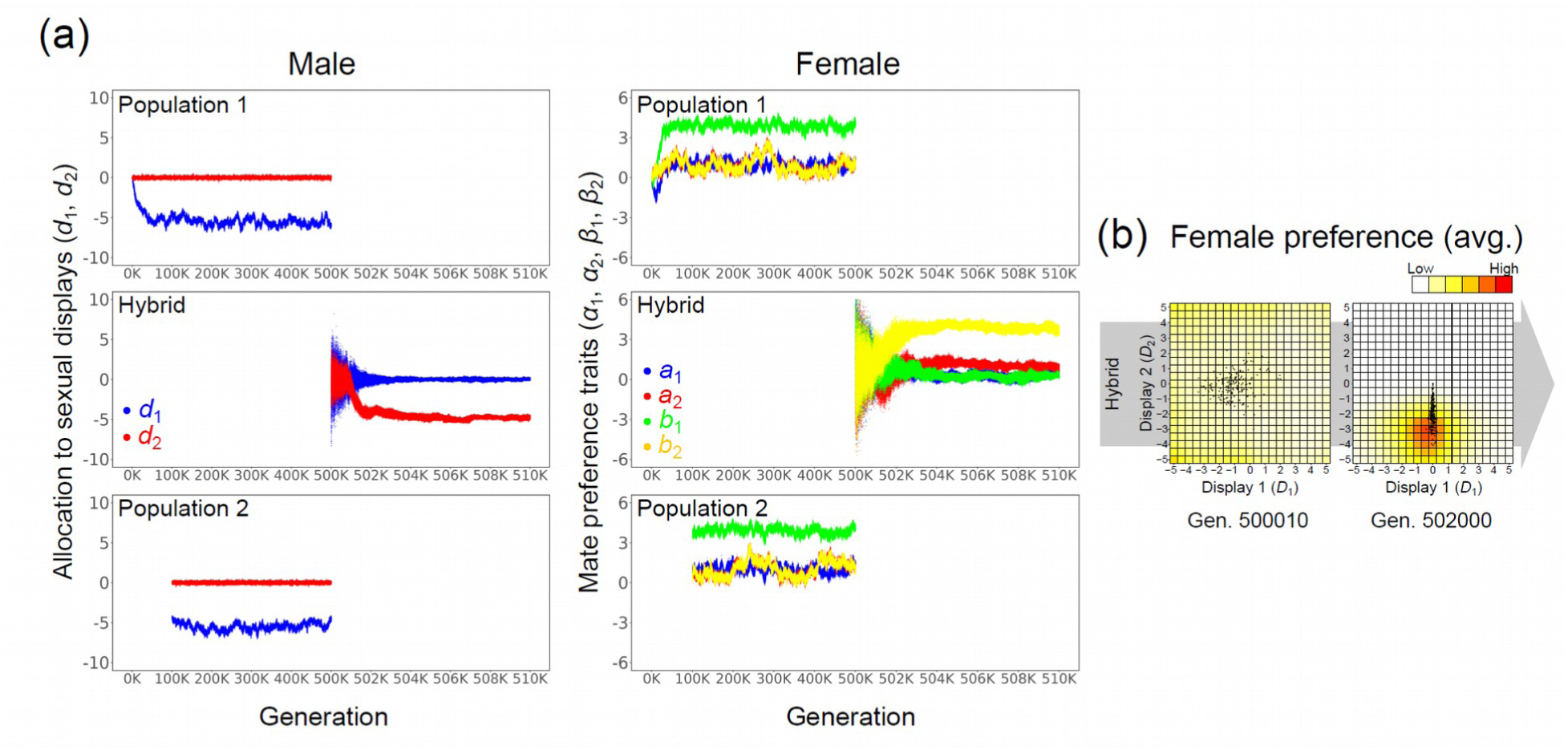
An example of hybrid speciation through one-time immigration of parental lineages to the secondary contact zone in the simulation assuming the beta preference function model. (a) Evolutionary dynamics of two male genetic traits (blue: *d*_1_, red: *d*_2_) and four female genetic traits (blue: *a*_1_, red: *a*_2_, green: *b*_1_, yellow: *b*_2_) are shown in the same format as Figure 2a; however, evolutionary dynamics in populations 1 and 2 after the secondary contact were not simulated and are not shown. (b) Averaged female preference function across all female individuals (red colors) and sexual display phenotypes of all male individuals (dark gray points) in the hybrid population are shown for two timepoints (generations 500010 and 502000) in the same format as Figure 2b. Parameters were set to the default values (Table S1).

**Figure S12.**
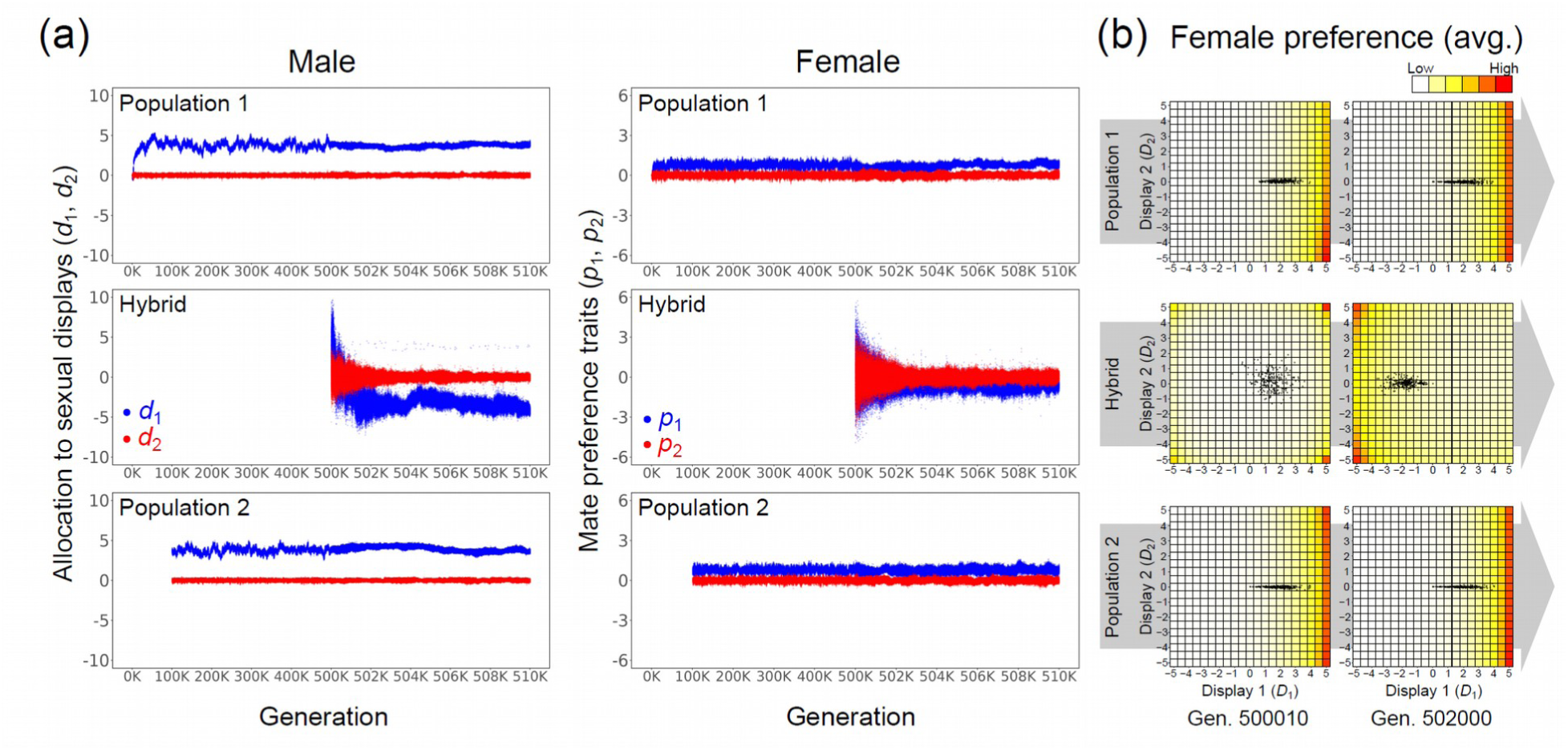
An example of hybrid speciation in the simulation assuming the exponential preference function model and the continuing immigration from parental-to the hybrid population. Evolutionary dynamics are shown in the same format as Figure 2 except that there are only two female traits (blue: *p*_1_, red: *p*_2_) in this simulation. Parameters were set to the default values (Table S1).

**Figure S13.**
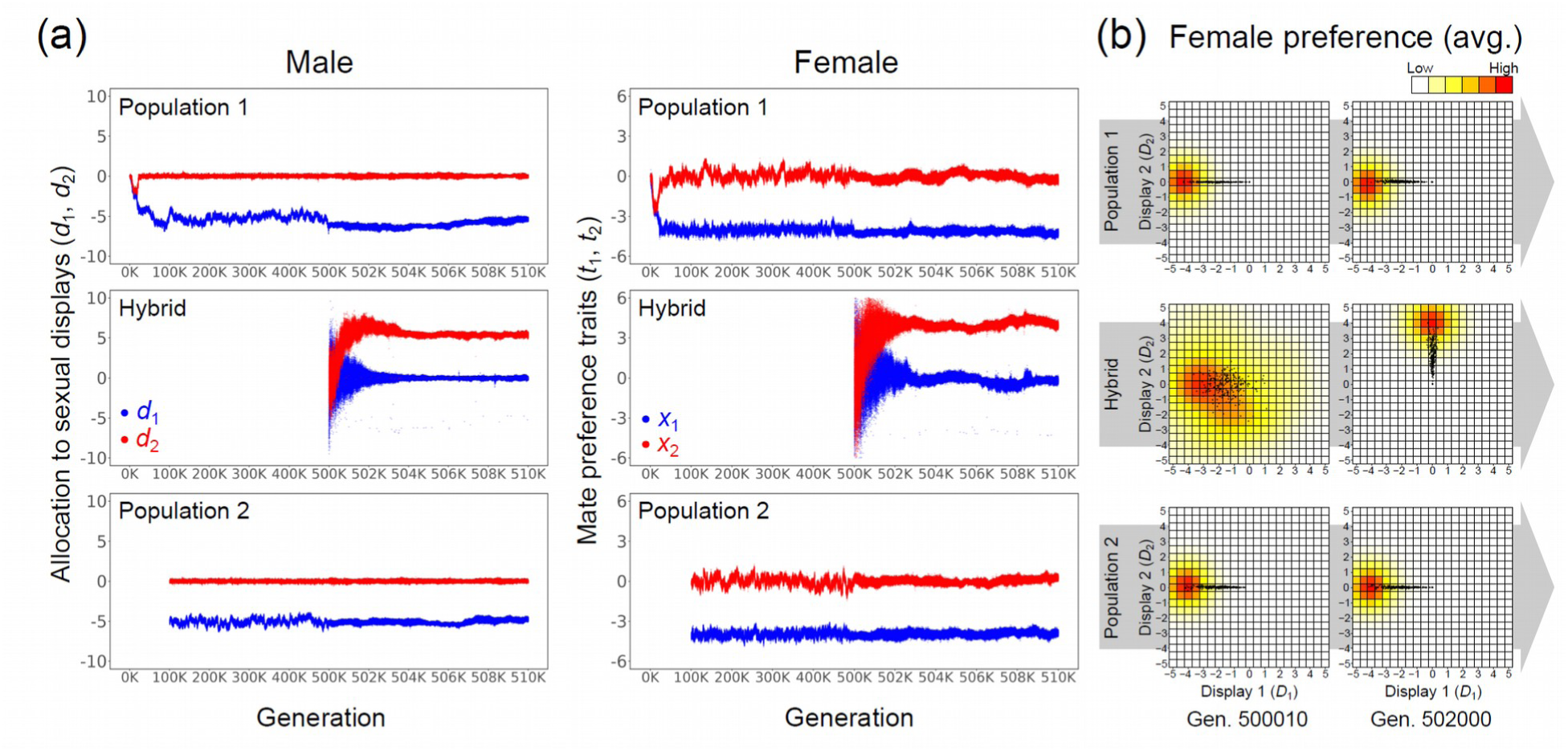
An example of hybrid speciation in the simulation assuming the Gaussian preference function model and the continuing immigration from parental-to the hybrid population. Evolutionary dynamics are shown in the same format as Figure 2 except that there are only two female traits (blue: *x*_1_, red: *x*_2_) in this simulation. Parameters were set to the default values (Table S1).

**Figure S14.**
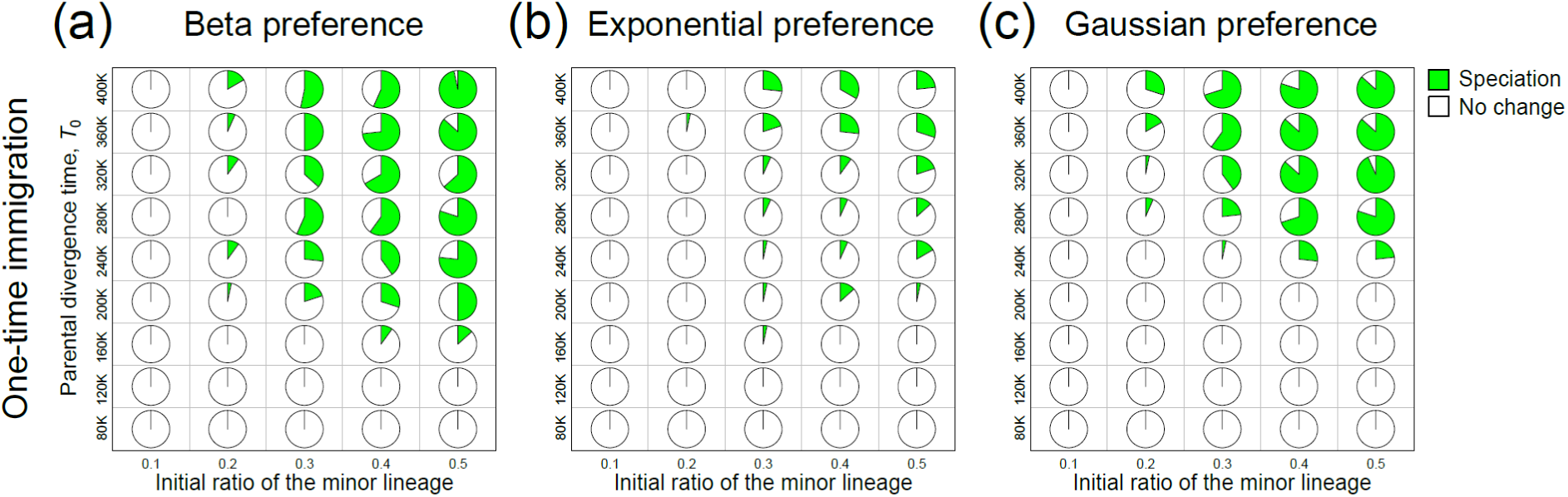
Conditions for hybrid speciation in simulations considering a secondary contact with “one-time immigration”, in which immigration from the parental populations to the contact zone occurs only once during the establishment of a population in the contact zone. Simulations were conducted with systematically varied values of *T*_0_ (the divergence time; vertical axis) and *r*_0_ (the initial ratio of two parental lineages in the hybrid population; horizontal axis). For each combination of parameter values, a pie chart shows frequencies of “speciation” (green) and “no change” (white) in 30 simulation replicates (other three categories of evolutionary outcome were not observed). Three panels show results of simulations with (a) the beta preference function model, (b) the exponential preference function model, and (c) the Gaussian preference function model, respectively. Other parameters were set to the default values (Table S1).

**Figure S15.**
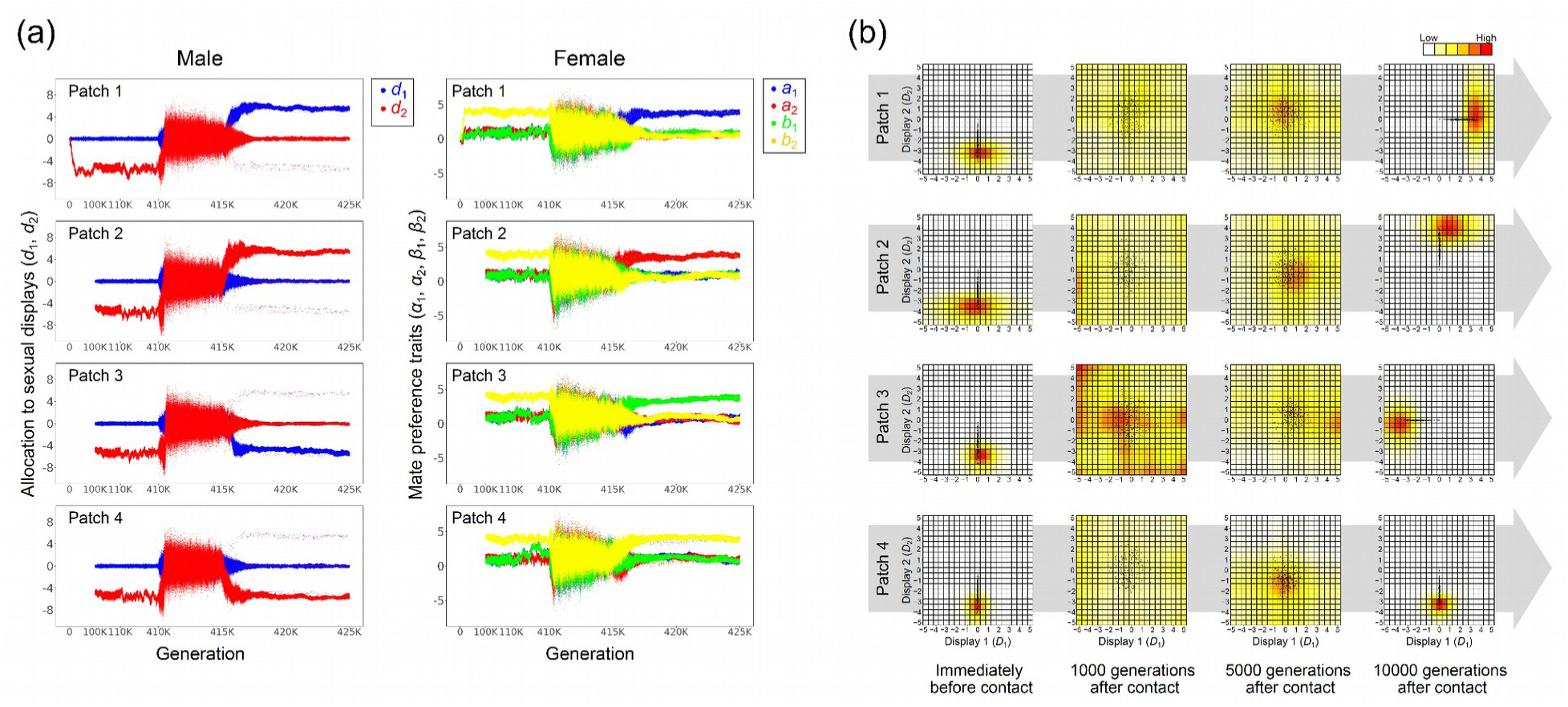
An example of simulation considering a cycle of isolation and reconnection of four habitable areas. (a) For each patch, values of two male genetic traits (blue: *d*_1_, red: *d*_2_) and four female genetic traits (blue: *a*_1_, red: *a*_2_, green: *b*_1_, yellow: *b*_2_) are shown for all male and female individuals at each generation. In the first 100K generations, the ancestral species evolved in a single population (patch 1), and the species spread to four patches in the following 10K generations (generations 100K - 110K). Then, an environmental change occurred in the generation 110K to cause the isolation of four patches (i.e. per-capita migration rate dropped to 0). After 30K generations (generation 410K), another environmental change occurred to cause the reconnection of four patches (i.e. migration rate was set to *κ* = 0.002), which led to a secondary contact of the four local populations. The secondary contact period (generations 410K - 425K) is magnified along the horizontal axis. (b) As in Fig. 1a, average mate preference function across all female individuals and sexual display phenotypes of all male individuals (dark gray points) are shown for each population at four time points (0, 1000, 5000, and 10000 generations after the onset of secondary contact at the generation 410K). Color of each cell shows the relative value of the preference score (i.e. the probability of acceptance) for each combination of male display phenotypes (*D*_1_, *D*_2_) to the score for the most preferred combination of male display phenotypes within the local population at that time point. Parameters were set to the default values (Table S1).

## Text S1: Details of the simulation model

Here I describe the following details of the simulations: three models of the mate preference function; procedures to simulate hybridization in a secondary contact zone; methods to identify stable equilibria of evolutionary dynamics; methods to quantify and classify simulation results.

### Preference function models

The model assumes that each female individual has a preference function, which maps the mating probability upon an encounter to every possible phenotype of male sexual displays (I did not adopt the “best-of-n” rule since the model will consider the mate search cost as the cost to be choosy). Most existing theoretical models of sexual selection considering continuous variation of mate preference use mathematically simple functions with a few parameters to express the preference function; genetic variation of preference function is considered by assuming such parameters as genetic traits. The way to model the preference function determines the range of possible shapes of the preference function as well as the relationship between genotypes and shapes of the preference function (e.g., the range of preference function shapes accessible from a given genotype by a few mutations). Evolutionary dynamics thus may strongly depend on the way to model the preference function. To evaluate the robustness of simulation results to changes in the preference function model, I conduct simulations with three alternative models of the preference function: the exponential-, Gaussian-, and beta preference function models (Fig. S1). Conventionally, many theoretical models of sexual selection used mathematically convenient functions such as the exponential or the Gaussian functions to formulate the preference function but both ways of modeling introduce hard artificial constraints in evolutionary dynamics [39]. For example, in models using the exponential function, mate preference is always open-ended, and preferences for certain male phenotypes cannot evolve even when female fitness will be maximized by choosing males with phenotypes in a certain range; in models using the Gaussian function, in contrast, evolution of open-ended preferences is not allowed. The beta preference function model partially solves these problems by allowing evolution of more flexible shapes of preference functions including unimodal, one-sided open-ended, and two-sided open-ended preferences. To my knowledge, one previous study [47] adopted a similar approach of using beta distribution function to model the mate preference function.

#### Exponential preference function model

This model considers two quantitative female traits, *p_s_*(*s* = 1, 2), each of which determines the direction and intensity of preference for the *s*-th display trait. Females with positive (negative) *p_s_*prefer males with positive (negative) phenotypic values of sexual display (*D_s_*) (Fig. S1a). Using an exponential function, the preference score from the female individual *j* to the *s*-th display of the male individual *i*, *P_sij_*, is given as: *P_sij_* = *exp*(*p_sj_*(*D_si_* − *D_s_best_*)), where the value of *D_s_best_* is the most preferred possible phenotype for the *s*-th display. The value of *D_s_best_* is *E_max_*if *p_sj_* ≥ 0 and −*E_max_* if *p_sj_*< 0. With this model, positive (negative) values of *p_s_* give a one-side open-ended preference function in favor of larger (smaller) values of *D_s_*, whereas *p_s_* = 0 yields random mating.

#### Gaussian preference function model

This model considers two quantitative female traits, *x_s_*(*s* = 1, 2), each of which corresponds to the target of preference for the *s*-th display trait (Fig. S1b). Using the Gaussian function, the preference score is given as follows: *P_sij_* = *exp* (−(*D_si_*− *x_sj_*)^2^).

#### Beta preference function model

This model uses the beta distribution function, which can represent various shapes with only two positive parameters *α* and *β*. Females have four quantitative traits, *a_s_* and *b_s_* (*s* = 1, 2) that determine the four parameters of the beta distribution function as: *α_s_* = *exp*(*a_s_*) and *β_s_*= *exp*(*b_s_*). Since the domain of the beta distribution function is 0 to 1, male display trait values is first mapped to this range using a logistic function as: *D_si_*\* = 1/(1+Σ*exp*(−*D_si_*)). Then, the preference score for a male individual *i*, whose sexual display is *D_si_*, by a female individual *j* is given as: *P_sij_* = *Beta*(*D_si_*\*; *α_sj_*, *β_sj_*) / *Beta*(*D_s_best_*\*; *α_sj_*, *β_sj_*), where *Beta*(*x*; *α*, *β*) represents the beta distribution function and *D_s_best_*\* is the value of the *s*-th display trait that gives the maximum value for *Beta*(*x*; *α_sj_*, *β_sj_*) within the range of possible values of *D_si_*\*. When *α* > 1 and *β* > 1, the beta distribution function has a finite maximum value at *x* = (*α* −1)/(*α* +Σ *β* − 2). Otherwise, beta distribution function diverges at *x* = 0 and/or *x* = 1; in this case, *D_s_best_*\* is 1/(1+Σ*exp*(−*E_max_*)) or 1/(1+Σ*exp*(*E_max_*)), the boundaries of the possible range of *D_si_*\*. Figure S1c shows example shapes of the beta preference function with several different combinations of female trait values.

### Details of the procedure for simulating scenario 1

Hybridization between two allopatrically diverged lineages in their secondary contact zone is simulated with the following procedure. First, the evolution of a common ancestral population is simulated for 100,000 generated, starting from the initial population of 100 individuals with the genome with no mutation, in order to eliminate influences of the artificial initial state. Second, the common ancestral population is divided into two isolated populations, and evolution of two populations is simulated for 500,000 generations, during which genomes of all individuals are recorded once per 10,000 generations. This procedure is repeated for 30 times in order to generate replications of parental lineage pairs for replicating the simulation of evolutionary dynamics in a secondary contact zone. Then, evolution in a secondary contact zone between two parental populations is simulated with various values of divergence time, *T*_0_ (*T*_0_ ≤ 500,000). Prior to starting a simulation of a secondary contact, two parental populations at the time point of *T*_0_ generations after their isolation were reconstructed from the record of genotypes of all individuals at that time point. Finally, a secondary contact between two populations is simulated by considering the third site (i.e., the contact zone) receiving immigration from both populations.

Immigration from two parental populations to the contact zone occurs in either “continuous immigration mode” or “one-time immigration mode”. With continuous immigration, the contact zone originally does not harbor a population, and immigration of individuals from both parental populations to the contact zone occurs recurrently. Every generation, the number of immigrants is given as a random number drawn from a Poisson distribution with intensity *m*. With one-time immigration, on the other hand, immigration from the parental populations to the contact zone occurs only once during the establishment of a population in the contact zone by 20×(1 − *r*_0_) individuals from the parental population 1 and 20×*r*_0_ individuals from the parental population 2. Parameter *r*_0_ is the ratio of two lineages in the founders of the hybrid population, which is applicable only to the one-time immigration mode. Both immigration modes do not consider immigration from the contact zone to parental populations for simplicity of the interpretation of simulation results.

### Methods to identify stable evolutionary equilibria

Conditions for the evolution of exaggerated sexual displays and the existence of multiple stable evolutionary equilibria are explored by simulating evolutionary dynamics of a closed population, starting from the initial population with 100 individuals with the genome containing no mutations. With the genome with no mutations, males have the most cryptic sexual display phenotypes, and females have a preference function without any bias toward specific exaggerated sexual displays under all three preference function models. Stable equilibrium of the evolutionary dynamics is identified based on the evolutionary dynamics of the mean values of two male genetic traits across all males in the population, ⟨***d***⟩ = (⟨*d*_1_⟩, ⟨*d*_2_⟩), which is recorded once per 100 generations. The two-dimensional space of two male genetic traits (*d*_1_, *d*_2_) is divided into five areas (Fig. 1a; hereafter, “trait-areas”), and if the mean male trait ⟨***d***⟩ stayed in the same trait-area for more than 10,0000 generations, the population was judged to be in an evolutionary stable equilibrium. The first area, area 0, corresponds to cryptic sexual displays; if a male’s genetic traits are in the area 0, resource allocation to sexual displays results in less than 10% reduction of offspring fitness when his resource budget size equals the expected value (*E_i_*= *E_max_*/2). Remaining part of the trait space is divided into four areas by two lines: *d*_2_ = *d*_1_ and *d*_2_ = − *d*_1_ (Fig. 1a). These four areas (areas 1, 2, 3, and 4) correspond to costly exaggerated sexual displays.

### Categorization of evolutionary outcomes in simulations of scenario 1

Evolutionary outcomes with the simulation scenario 1 are sorted into five categories to examine frequencies of speciation and other outcomes for each parameter setting. The categorization is based on (i) whether or not mean values of male genetic traits in the hybrid population, ⟨***d***⟩ = (⟨*d*_1_⟩, ⟨*d*_2_⟩), has reached an evolutionary stasis and (ii) the strength of the barrier to gene flow (*B*) between the hybrid population and parental populations in the final 100 generations of the simulation. Male genetic traits are judged to have reached an evolutionary stasis if ⟨***d***⟩ has stayed in the same area of the trait space (“trait-area” shown in the Fig. 1a) throughout the last 5000 generations of the secondary contact period. This judgment is based on records of mean male traits with the frequency of once per 100 generations. In cases with continuing immigration, *B* is measured as 1 − *R_i_*/*R_r_*, where *R_i_* is the average number of offspring produced by immigrant individuals through mating with resident individuals, and *R_r_*is the average number of offspring produced by resident individuals through mating with resident individuals. In the absence of premating isolation, *R_i_*and *R_r_* are expected to be equal, whereas in the presence of complete premating isolation, *R_i_* should approach zero. I first calculate *B* between immigrant male and resident female and *B* between immigrant female and resident male separately. Then, the final score of *B* is given as the average value of them. In simulations with one-time immigration, where the above formula is not applicable, the value of *B* is always set to 1. Using these statistics, evolutionary outcomes of hybridization are divided into the following categories: (1) “speciation”, where *B* ≥ 0.8 and ⟨***d***⟩ has reached a stasis in a trait-area that is different from that of the parental populations and trait-area 0; (2) “incomplete speciation”, where ⟨***d***⟩ has reached a stasis in a trait-area that is different from that of the parental populations and trait-area 0, but *B* < 0.8; (3) “no change”, where ⟨***d***⟩ has reached a stasis in the trait-area same as that of the parental populations; (4) “cryptic display”, where ⟨***d***⟩ has reached a stasis in the trait-area 0; (5) “drift”, where ⟨***d***⟩ has not reached a stasis.

### Methods to count the number of species in simulations of scenario 2

I counted the number of species with distinct exaggerated sexual displays that have been formed within 10,000 generations after the start of the secondary contact period. For this purpose, the mean male trait ⟨***d***⟩ in each local population was tracked in the period from 10,000 to 15,000 generations after the start of secondary contact in order to determine whether and in which trait-area (Fig. 1a) the mean male trait has reached an evolutionary stasis. The number of species with different exaggerated sexual displays is measured by counting the number of trait-areas (areas 1, 2, 3, and 4) within which one or more local populations have reached stasis. Local populations that have not stayed in a single trait-area or have stayed in the area 0 are not counted as species with exaggerated sexual displays. Local populations that have stayed in the same trait-area are counted together as a single species.

## Notes

### Competing Interest Statement

The authors have declared no competing interest.

